# Comparative transcriptomics reveal highly conserved regional programs between porcine and human colonic enteric nervous system

**DOI:** 10.1101/2022.02.24.480770

**Authors:** Tao Li, Marco Morselli, Trent Su, Million Mulugeta, Muriel Larauche, Matteo Pellegrini, Yvette Taché, Pu-Qing Yuan

## Abstract

The porcine gut is increasingly regarded as a useful translational model. The enteric nerve system (ENS) in the colon coordinates diverse functions. However, knowledge of the molecular profiling of porcine ENS and its similarity to that of human is limited. We identified the distinct transcriptional programs associated with functional characteristics between inner submucosal and myenteric ganglia (ISG, MG) in porcine proximal and distal colon (p-pC, p-dC) using bulk RNA sequencing (RNA-seq) and single-cell RNA-seq. Comparative transcriptomics of MG in corresponding colonic regions of porcine and human revealed highly conserved programs existing in p-pC and p-dC, which explained >90% of their transcriptomic responses to vagal nerve stimulation (VNS), suggesting that p-pC and p-dC could serve as predictors in translational studies. The conserved programs specific for inflammatory modulation were displayed in porcine with VNS. This study provides a valuable transcriptomic resource for understanding of human colonic functions and neuromodulation using porcine model.

## Introduction

The porcine gastrointestinal (GI) tract is increasingly regarded as a useful translational research model due to its structural and functional similarities with human. Compared to mouse, rat, dog, cat or horse, both porcine and human are omnivores and colon fermenters, and have similar metabolic and intestinal physiological processes and microbial composition^1–4^. Among the important similarities relevant to colonic function, both taenia and haustra/sacculation are present in porcine and human colon, while missing in dogs and most rodents, despite the porcine proximal colon being orientated in a spiral fashion^2,5^. Additionally, compared with human, porcine has a similar chromosomal structure and a comparably sized genome with a 60% sequence homology and 93% correspondence in relevant biomarkers^6,7,8^. The porcine size is also relevant closely mimicking drug dose volumes used in human and for evaluating medical devices^9^. Thus, the porcine model has untapped potential for a greater understanding of the underlying colonic functions of human in health and diseases ^10^.

The GI tract is highly innervated by both extrinsic (sympathetic and parasympathetic) and intrinsic (enteric nervous system, ENS) nervous systems. The ENS is endowed with complex reflex circuits that control a variety of GI physiological functions via networks of neurons and glia interacting with cells located in the mucosal and muscular layers^11^. Uniquely the ENS is the only part of the peripheral nervous system that can function independently of central nervous system innervation. Therefore, an intact ENS is essential for life and its dysfunction is often linked to digestive disorders^11,12^. In mammalian colon, the ENS comprises two main ganglionated plexuses: myenteric plexus, located between longitudinal and circular layers of muscularis externa, and submucosal plexuses (SMP) within the submucosa. Unlike the SMP in rodents that is single-layered^13^, both porcine and human colonic SMP consists of an inner submucosal plexus, located close to muscularis mucosae, and an outer submucosal plexus, located on the luminal side of the circular muscle layer^14^. Despite the anatomical similarity, it is still unknown to what extent transcriptional programs are conserved between porcine and human colonic ENS. Such information is of importance for the use of porcine model in translational studies.

Recent studies in mice have demonstrated heterogeneity in neuronal identity, morphology, projection orientation and synaptic complexity within the ENS located in different colonic segments in line with the diversity of colonic motility patterns^15^. The region-dependent molecular characterization of porcine ENS has been little investigated so far. In addition, colonic functions are modulated through parasympathetic innervation, responsible for regulating colonic secretion and motility^16^. We have recently reported that electrical vagal nerve stimulation (VNS) triggered pan-colonic contractions in the porcine^17^. However, it is unknown how such VNS influences colonic transcriptional programs and whether the impacts of VNS are region-specific.

The incomplete ENS characterization is largely due to longstanding technical challenges in isolating enteric neurons^18^. Laser-capture microdissection (LCM) has been the method of choice for accurately targeting and capturing cells of interest from a heterogeneous tissue sample^19^ like the colon^20^. The enriched cells can be used for bulk RNA sequencing (RNA-seq), which becomes a widely adopted method for profiling transcriptomic variations^21^ and is a promising choice to characterize the region-dependent molecular profiling in ENS. More recently, single-cell RNA sequencing (scRNA-seq) has emerged as a powerful method for exploring gene expression profile at single-cell resolution^22^. However, its high cost limits its application across a large population of samples^23^. Considering this constraint, several computational methods have been exploited to deconvolve cell type composition in bulk RNA-seq data using scRNA-seq reference datasets^24–27^ which largely facilitate the use of scRNA-seq data from a small number of subjects in clinical setting involving a large number of subjects.

In the present study, we profiled the transcriptomes of myenteric and inner submucosal ganglia (MG, ISG) from porcine proximal, transverse and distal colon (p-pC, p-tC, p-dC), and MG in parallel from human ascending, transverse and descending colon (h-aC, h-tC, h-dC) using LCM coupled with bulk RNA-seq analysis. By cross-comparing the transcriptional profiling of MG in the corresponding colonic regions between porcine and human, we aimed to characterize the conservation of gene programs derived from orthologous genes. Then, based on the high-quality orthologous genes, we evaluated discrepancy of the region-specific gene programs in MG and ISG after electrical stimulation of the celiac branch of the abdominal vagus nerve in anesthetized porcine. Additionally, we performed scRNA-seq coupled with bulk gene expression deconvolution to identify cell-type marker genes and putative interactions among neurons and glia in MG of porcine colon and further revealed the regional transcriptomic heterogeneity in the ENS of porcine with and without VNS.

## Results

### Transcriptomic similarities of colonic myenteric ganglia between porcine and human

The obtained empirical cumulative density function (ECDF) profiles and both Kolmogorov-Smirnov and Pearson’s Chi-squared test (*p* > 0.05) based on the averaging normalized transcription levels of each orthologous gene demonstrated that gene expression profiling in MG of human colonic segments (h-aC, h-tC, h-dC) could be predicted according to that of corresponding porcine colonic segments (p-pC, p-tC, p-dC), following the probability distribution shown in ECDF profiles (Fig. 1a, c). In addition, pair-wise comparison addressed a significant Spearman’s correlation (*p* < 0.001) (Fig. 1b).

**Fig. 1.**
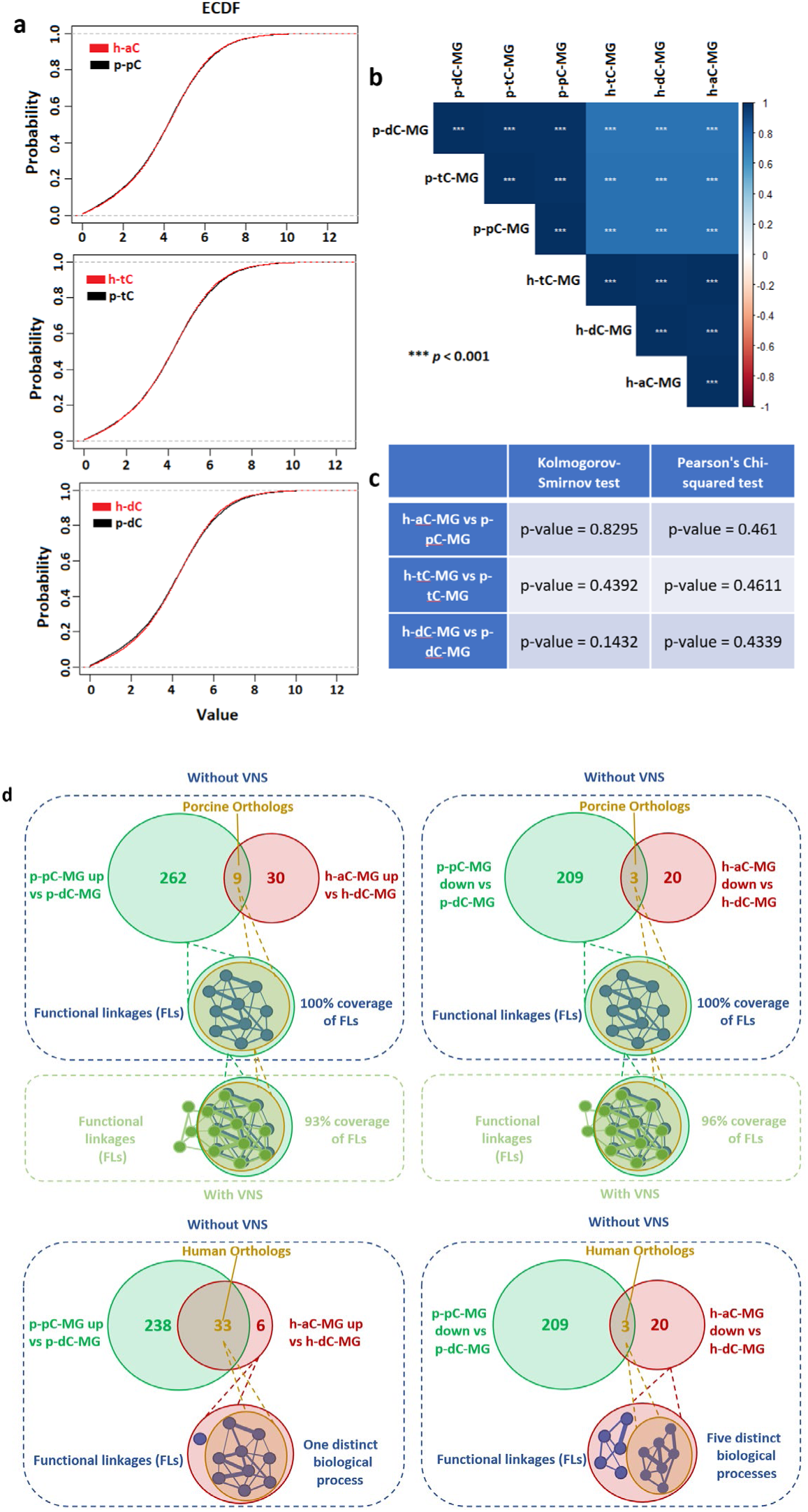
Probability distributions of human-porcine regional datasets selected from the high-quality orthologous gene list between myenteric ganglia (MG) from porcine proximal, transverse and distal colon (p-pC, p-tC, p-dC) and in parallel from human ascending, transverse and descending colon (h-aC, h-tC, h-dC) (a), Spearman’s Correlation (b) and evaluation of goodness of fit (c). Cartoon diagram showing regional differentially expressed gene (DEG) network matching between human and porcine. FLs, functional linkages (d). VNS, vagal nerve stimulation. Up, upregulated expression. Down, downregulated expression.

The differentially expressed genes (DEG) enriched significantly (*p* < 0.05) by Gene Ontology terms were selected for comparisons among colonic segments, between MG and ISG, and the porcine and human. The functional linkages related to the orthologous DEG were extracted to reflect functional similarities across species. Twelve porcine orthologous DEG (9 upregulated and 3 downregulated) in comparison of h-aC-MG and h-dC-MG were involved in the regulatory networks, mediated by 483 DEG (271 upregulated and 212 downregulated), when comparing p-pC-MG and p-dC-MG (Fig. 1d). The functional linkages related to the 12 porcine orthologous DEG which achieved 100% linkage coverage and represented all the characteristics of p-pC-MG and p-dC-MG. Moreover, these conserved functional linkages accounted for more than 90% of all functional linkages of the regulatory networks responding to VNS when comparing p-pC-MG and p-dC-MG. However, partial coverage was shown when matching the functional linkages derived from human orthologous DEG in comparison of p-pC-MG and p-dC-MG and DEG in comparison of h-aC-MG and h-dC-MG (Fig. 1d). We did not find any porcine orthologous DEG when comparing h-aC-MG or h-dC-MG with h-tC-MG. Therefore, only p-pC and p-dC were considered in the subsequent analyses.

### Preferential enrichment for genes associated with MG and ISG functions in porcine proximal or distal colon

The functional linkages related to each DEG list in our study and those related to the DEG list screened by the high-quality human orthologous genes achieved almost 100% linkage coverage and the cell-subtype gene markers also fell into the high-quality orthologous gene list. Two databases (Gene Ontology Biological Processes and WikiPathways) were used to interpret the RNA-seq data. The interactions were defined when the networks mediated by DEG involved in WikiPathways of interest and Biological Processes (BPs) shared BPs. We mainly focused on the scenario where there were interactions between the pathways of interest unless otherwise stated.

Innate similarities between p-pC and p-dC were found when comparing respective MG and ISG. The upregulated genes in MG of both p-pC and p-dC accounted for a larger percentage of identified genes involved in the top five BPs and led to the higher enrichment of the BPs (Supplementary Fig. 1). The BPs connected with those such as nerve development, neuronal projection guidance and neuron/glial cell differentiation. In addition, these genes in MG contributed to higher enrichment scores of the WikiPathways such as oncostatin M signaling pathway, glutamate binding, activation of AMPA receptors and synaptic plasticity, chemokine signaling pathway, nitric oxide biosynthesis, dopaminergic neurogenesis and TGF-beta receptor signaling, while the downregulated genes in MG of both p-pC and p-dC made contribution to higher enrichment scores of the WikiPathways such as activation of angiotensin pathway, acetylcholine synthesis, synaptic vesicle pathway, neurotransmitter release cycle and dopamine transport and secretion (Fig. 2).

**Fig. 2.**
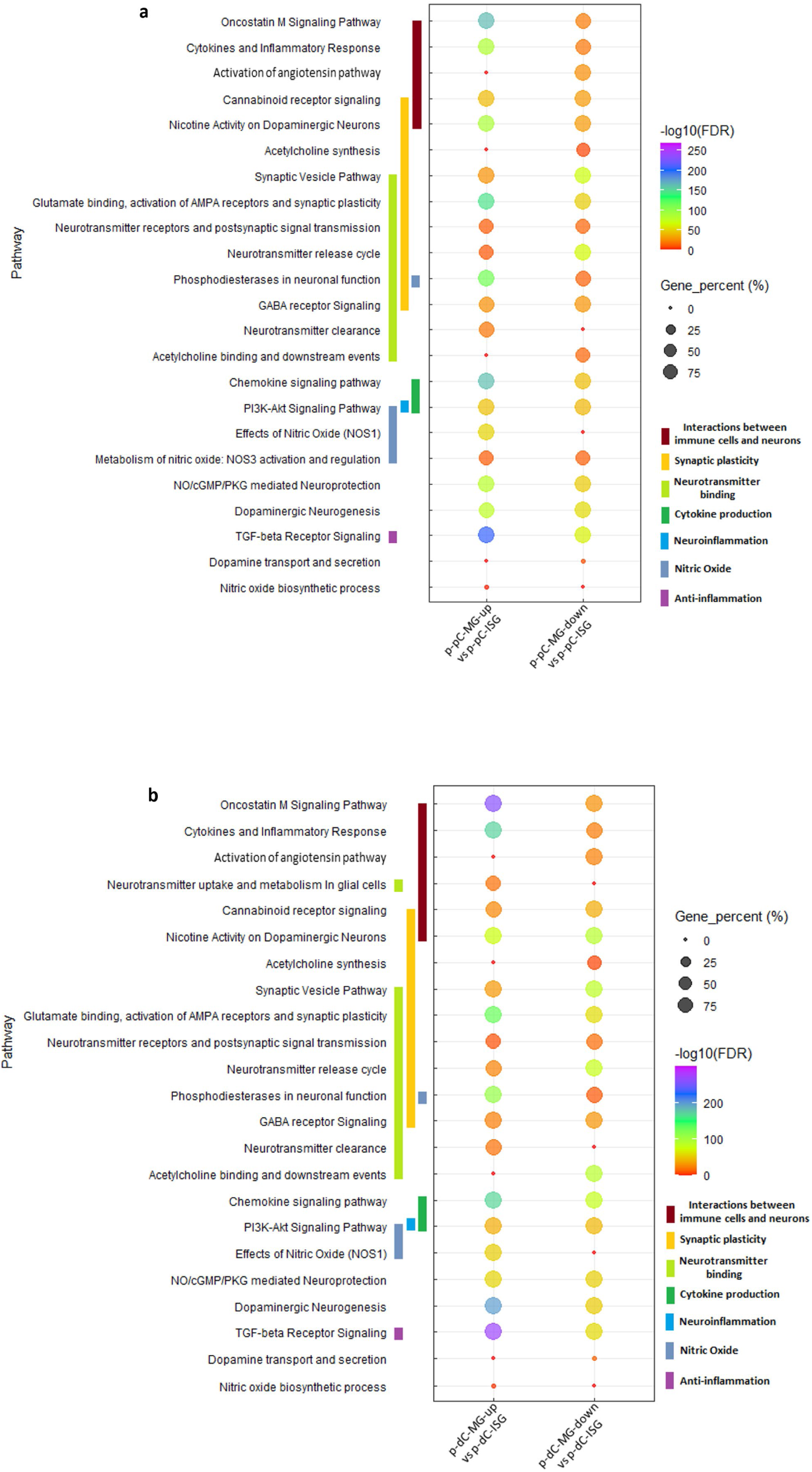
Comparison of pathway enrichment between myenteric ganglia (MG) and inner submucosal ganglia (ISG) in porcine proximal (a) or distal colon (p-dC, p-dC) (b). The bubble plots show the gene percentage and enrichment of WikiPathways. Circle size represents the ratio of the number of pathway-specific differentially expressed genes (DEG) and the number of total DEG in each DEG list (Gene_percent). Color represents a -log10(FDR) distribution from big (orange) to small (purple). The color bars mean the specified WikiPathways classified into designated clusters. Up, upregulated expression. Down, downregulated expression.

However, there were still some differences found in comparison of MG and ISG between p-pC and p-dC. The upregulated genes in p-pC-MG contributed to higher enrichment scores of the WikiPathways such as nicotine activity on dopaminergic neurons, while the upregulated genes in p-dC-MG exclusively resulted in neurotransmitter uptake and metabolism in glial cells (Fig. 2).

### Pathway enrichment changes in MG or ISG in the proximal-distal axis of the porcine colon

In p-pC-MG, the downregulated genes contributed to larger enrichment scores of the BPs (Supplementary Fig. 2a, c), which connected with ENS development, blood vessel development and glial cell differentiation, while the BPs involving the upregulated genes connected with ENS development, synaptic organization and chemical synaptic transmission. In ISG, the enrichment differences between p-pC and p-dC were not as large as those between p-pC-MG and p-dC-MG. The biggest enrichment disparity happened to blood vessel development and the downregulated genes in p-pC-ISG contributed to its higher enrichment score (Supplementary Fig. 2b, d).

The upregulated genes in p-pC-MG led to larger enrichment scores of the WikiPathways such as acetylcholine synthesis, synaptic vesicle pathway, neurotransmitter receptors and postsynaptic signal transmission, neurotransmitter release cycle and dopamine uptake, unlike those relate to Oncostatin M signaling pathway, cytokines and inflammatory response, chemokine signaling pathway, NO/cGMP/PKG mediated neuroprotection, dopaminergic neurogenesis, TGF-beta receptor signaling and smooth muscle contraction (Fig. 3a). All listed pathways interacted with smooth muscle contraction. The upregulated genes in p-pC-ISG contributed to most prominent enrichment scores of nicotine activity on dopaminergic neurons, neurotransmitter receptors and postsynaptic signal transmission, dopaminergic neurogenesis, and dopamine biosynthesis process. The downregulated genes in p-pC-ISG resulted in enrichment scores related to nitric oxide biological action, chemokine signaling pathway and TGF-beta receptor signaling (Fig. 3b).

**Fig. 3.**
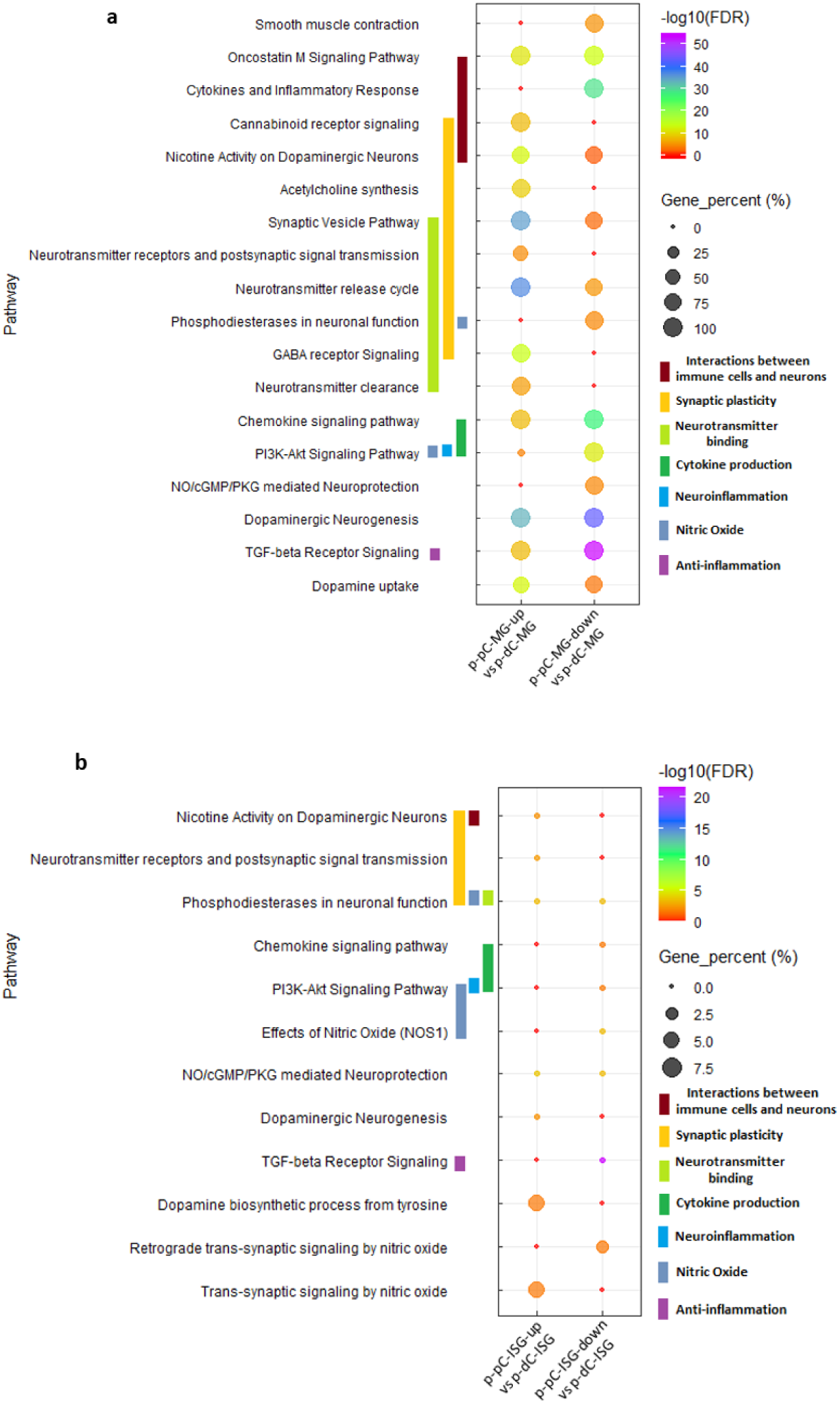
Comparison of pathway enrichment in myenteric ganglia (MG) (a) or in inner submucosal ganglia (ISG) (b) between porcine proximal and distal colon (p-pC, p-dC). The bubble plots show the gene percentage and enrichment of WikiPathways. Circle size represents the ratio of the number of pathway-specific differentially expressed genes (DEG) and the number of total DEG in each DEG list (Gene_percent). Color represents a -log10(FDR) distribution from big (orange) to small (purple). The color bars mean the specified WikiPathways classified into designated clusters. Up, upregulated expression. Down, downregulated expression.

### Distinct cell-type-specific pathway enrichment across porcine colonic segments detected by scRNA-seq

The scRNA-seq data showed that there were 10 cell clusters in muscularis externa of p-pC, p-tC (data not shown) and p-dC detected with similar cell markers (Fig. 4a, b). The neuronal and glial clusters were annotated with markers *GAP43* and *CLDN11*, respectively, which were validated by single-molecule fluorescence hybridization (smFISH). *GAP43* and *CLDN11* were co-expressed with *ELAVL4* and *GFAP*, respectively (Fig. 5A-C, M-O). Cell-cell interaction analysis revealed the strongest neuron-glia interaction in p-pC, while the interactions in p-tC (data not shown) and p-dC were almost the same (Fig. 4a, b). Three neuronal and two glial sub-clusters were further annotated with markers *ACLY* for cholinergic neurons, *GLS* for glutamatergic neurons, *ARHGAP18* for nitrergic neurons, *SLC41A1* and *SPC24* for two glial cell types and validated by smFISH with *ChAT*, *VGLU2*, *NOS1* and *GFAP*, respectively (Fig. 4c, d, 5D-L, P-S). Cell-cell interactions were also detected from ligand-to receptor-expressing neuronal and glial subsets including interactions between cholinergic neurons and nitrergic neurons or SLC41A1_ or SPC24_glia, glutamatergic neurons and cholinergic neurons or SLC41A1_glia, nitrergic neurons and SLC41A1_ or SPC24_glia, and SLC41A1_glia and SPC24_glia (Fig. 4e, Supplementary Table 1).

**Fig. 4.**
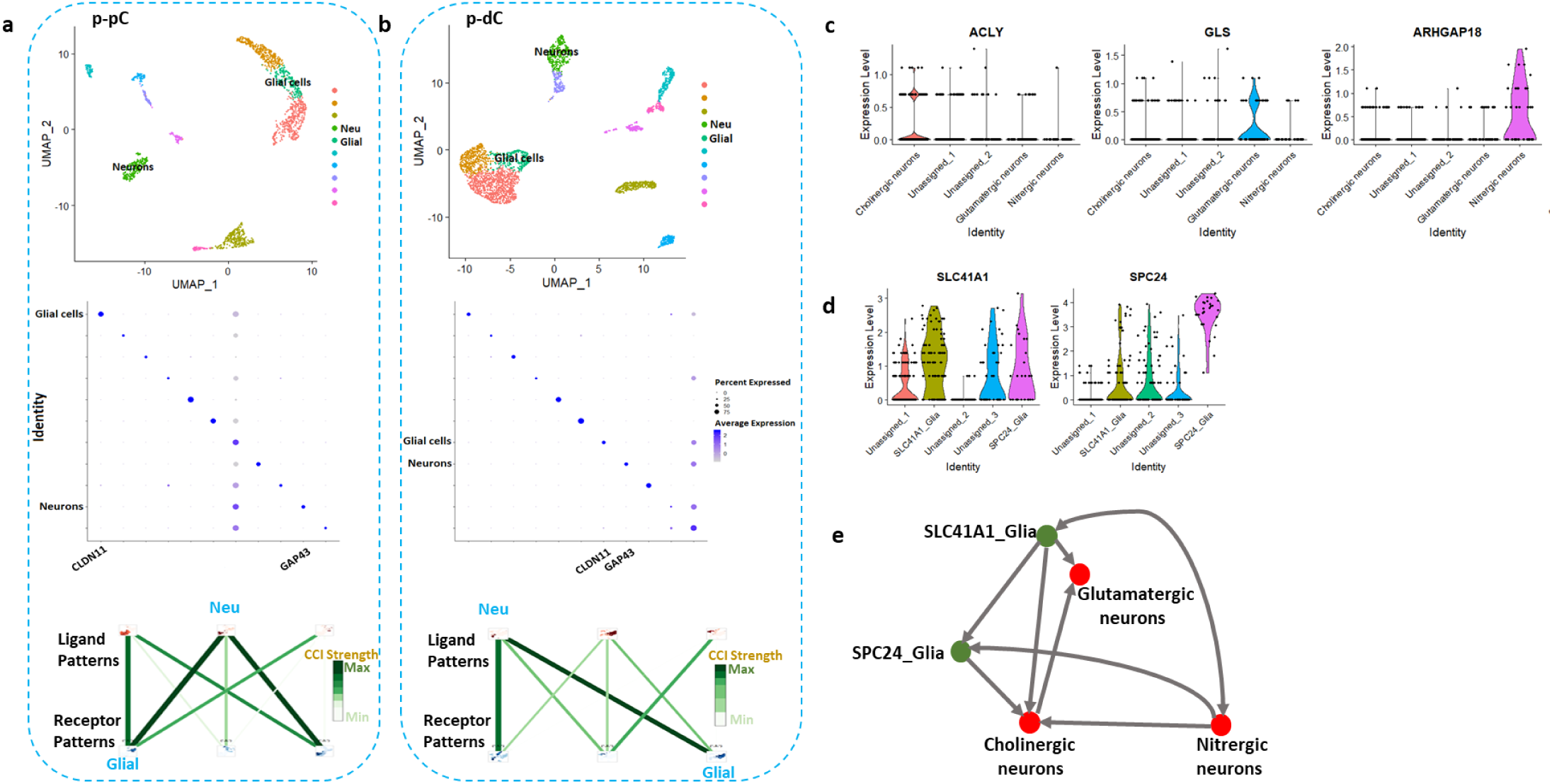
Visualization of cell types and their interaction strength in the muscularis externa of proximal colon (p-pC) (a) and distal colon (p-dC) (b) in naïve porcine by single-cell RNA sequencing. The clusters were labeled if the corresponding marked were verified by in situ hybridization. Uniform Manifold Approximation and Projection (UMAP) visualized 10 distinct cell clusters and neuronal (Neu), glial types and their interaction strength were highlighted only. Violin plots depict the distribution of gene expression levels for 3 neuronal subpopulations (c) and 2 glial subpopulations (d). Interactions (arrows) from ligand- to receptor-expressing neuronal and glial subsets (e). Red, neuronal subset; green, glial subset. Growth associated protein 43 (GAP43), a marker for annotating pan-neurons; Claudin 11 (CLDN11) for pan-glial cells; ATP citrate lyase (ACLY) for cholinergic neurons; Glutaminase (GLS) for glutamatergic neurons; Rho GTPase activating protein 18 (ARHGAP18) for nitrergic neurons; Solute carrier family 41 member 1 (SLC41A1) and SPC24 component of NDC80 kinetochore complex (SPC24) for two subtypes of glia.

**Fig. 5.**
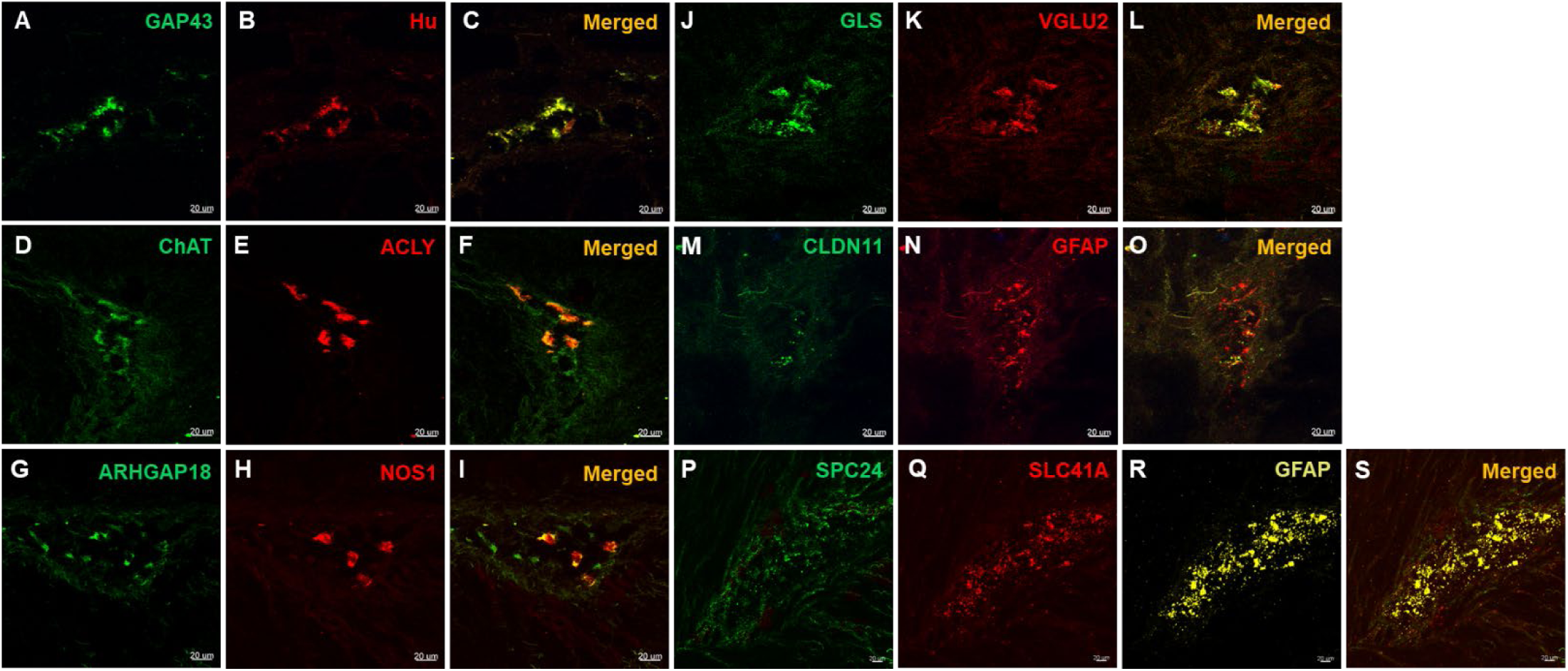
Validation of discriminatory marker genes for major classes of enteric neurons and two subtypes of enteric glial cells (EGCs) in the myenteric ganglia of porcine proximal colon using RNAscope in situ hybridization. Confocal images were generated from 5-10 optical sections (Z-stack) cross the myenteric ganglia with frame 436×436 μm and 1 μm apart (20× objective). The co-expression of mRNA for GAP43 (annotating enteric neurons) with Hu (A, B), ACLY (annotating cholinergic neurons) with ChAT (D, E), ARHGAP18 (annotating nitrergic neurons) with NOS1(G, H), GLS (annotating glutamatergic neurons) with VGLU2 (J, K), CLDN11 (annotating EGCs) with GFAP (M, O), and SPC24, SLC41A (annotating two subtypes of EGCs) with GFAP (P, Q, R) were indicated in the merged images (C, F, I, L, O and S) respectively.

The WikiPathways with top ranking enrichment scores in comparison of p-pC-MG and p-dC-MG were selected to evaluate contribution of each cell subset to pathway enrichment. Among the pathways involving the upregulated genes in p-pC-MG, cholinergic neurons played a leading role and contributed to enrichment of neurotransmitter release cycle, which was also attributed to glutamatergic neurons, enrichment of synaptic vesicle pathway, which was also attributed to SLC41A1_glia, and enrichment of chemokine signaling pathway and TGF-beta receptor signaling (Fig. 6). Among the pathways involving the downregulated genes in p-pC-MG, nitrergic neurons contributed to enrichment of dopaminergic neurogenesis, TGF-beta receptor signaling and chemokine signaling pathway. However, glutamatergic neurons contributed more to enrichment of chemokine signaling pathway. Although cholinergic neurons contributed to enrichment of neurotransmitter release cycle, their contribution was much less than that to enrichment of the pathway involving the upregulated genes in p-pC-MG (Fig. 6).

**Fig. 6.**
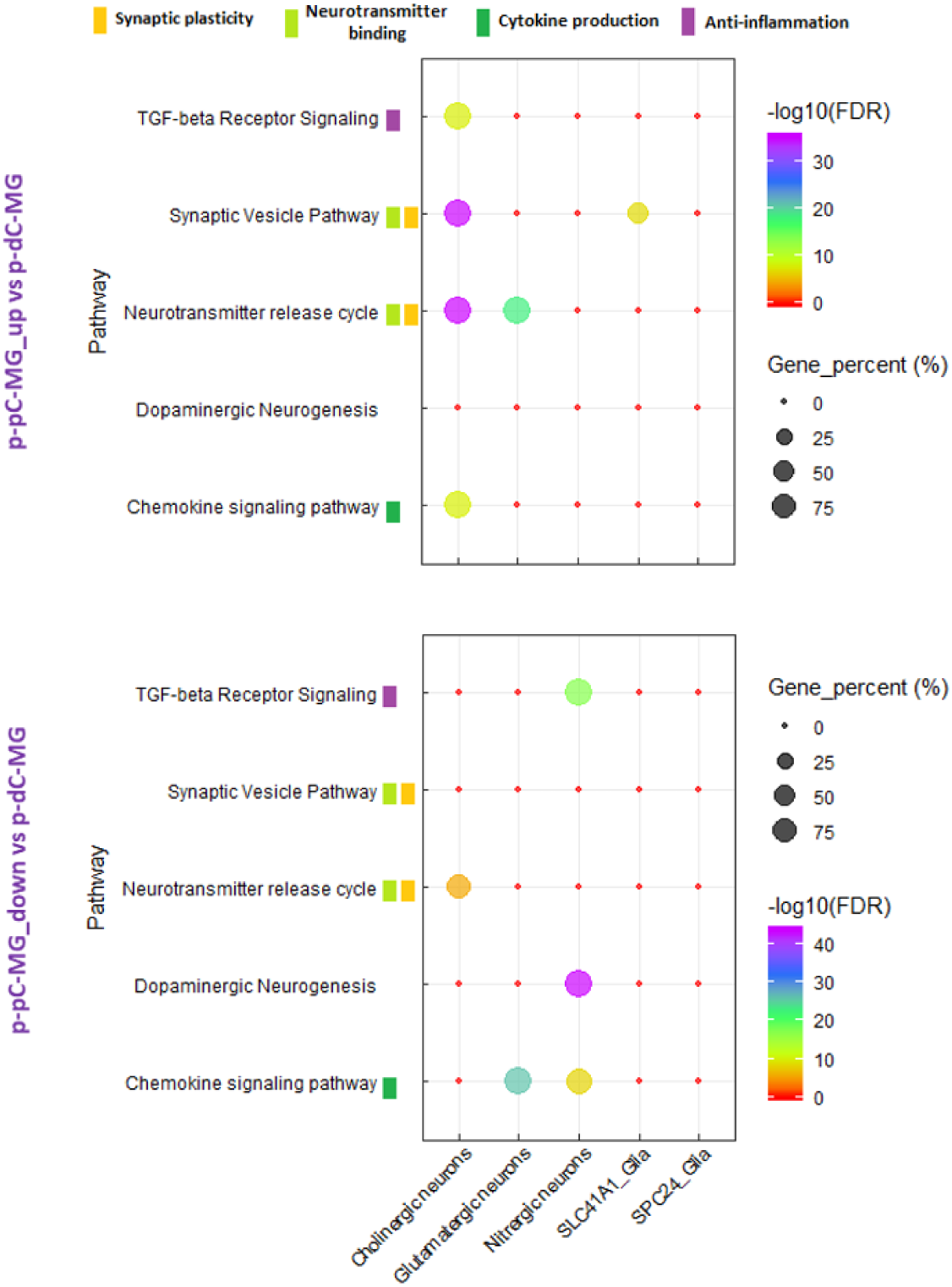
Comparison of cell-type specific pathway enrichment in myenteric ganglia (MG) between porcine proximal and distal colon (p-pC, p-dC). The bubble plot shows the gene percentage and enrichment of WikiPathways. Circle size represents the ratio of the number of cell-type pathway-specific differentially expressed genes (DEG) and the number of total DEG in each DEG list (Gene_percent). Color represents a -log10(FDR) distribution from big (orange) to small (purple). The color bars mean the specified WikiPathway classified into designated clusters. Up, upregulated expression. Down, downregulated expression.

### Region-specific effects of vagal nerve stimulation on pathway enrichment in porcine proximal MG and ISG and in porcine proximal and distal MG

The enrichment ratio of the same pathway involving upregulated and downregulated genes in each DEG list (e.g., p-pC-MG versus p-dC-MG, or p-pC-MG versus p-pC-ISG) was calculated and compared between porcine with and without VNS to evaluate its influences at both single and bulk resolution. The DEG, derived from comparison of p-dC-MG and p-dC-ISG in porcine with or without VNS, were involved in almost the same WikiPathways, despite different numbers of DEG. Only one differentially regulated gene under VNS was found, but enrichment of the WikiPathway mediated by the gene was not significant (Supplementary Fig. 4a), suggesting a significant overlap in the transcriptional landscape under these two situations. Similarly, comparison of p-pC-ISG and p-dC-ISG revealed that no differences were observed in porcine with and without VNS (Supplementary Fig. 4b).

Compared with porcine without VNS, BP enrichment patterns in porcine with VNS were almost the same based on comparison of p-pC-MG and p-pC-ISG. There were higher enrichment scores of the BPs involving upregulated genes in p-pC-MG than those involving downregulated genes in p-pC-MG. However, there were some alterations of BP enrichment patterns in porcine with VNS based on comparison of p-pC-MG and p-dC-MG. Enrichment scores of the BPs such as G protein-coupled receptor signaling pathway, anatomical structure morphogenesis and system process, involving upregulated genes in p-pC-MG, were higher than those involving downregulated genes in p-pC-MG (Supplementary Fig. 1, 2, 3). These BPs were connected with regulation of signaling, cell communication, immune response and immune system process regulated by VNS.

Based on comparison of p-pC-MG and p-dC-MG, VNS promoted enrichment of most of the WikiPathways in p-pC-MG, including gap junctions, pro- and anti-inflammatory signaling, chemokine signaling pathway, neurotransmitter uptake and metabolism in glial cells, NO/cGMP/PKG mediated neuroprotection, dopaminergic neurogenesis, nicotine activity on dopaminergic neurons, oncostatin M signaling pathway and TGF-beta receptor signaling, and reduced enrichment of human orthologous risk genes for intestinal and extra-intestinal diseases, synaptic vesicle pathway and neurotransmitter release cycle (Fig. 7a). By contrast, effects of VNS on p-dC-MG were reflected by improved enrichment of smooth muscle contraction, GABA receptor signaling and glutamate binding, activation of AMPA receptors and synaptic plasticity (Fig. 7a). In addition, the comparison between p-pC-MG and p-pC-ISG showed that VNS decreased the enrichment scores of most of the WikiPathways in p-pC-MG, such as gap junctions, anti-inflammatory signaling, smooth muscle contraction, chemokine signaling pathway, glutamate binding, activation of AMPA receptors and synaptic plasticity and TGF-beta receptor signaling, while increasing the enrichment scores of the WikiPathways, including pro-inflammatory signaling, acetylcholine binding and downstream events, GABA receptor signaling, neurotransmitter release cycle and synaptic vesicle pathway (Fig. 7b). The alterations were further explained by the functional similarity between gene products, which was measured by semantic similarities between the annotated BPs of each gene, resulting in similarity matrix (Supplementary Fig. 5). High average similarity scores in the WikiPathways of interest indicate strong functional similarities. Gap junctions and neurotransmitter release cycle had the highest similarity scores. High scores existed when comparing acetylcholine binding and downstream events or neurotransmitter release cycle with pro-inflammatory unlike anti-inflammatory signaling. Apart from functional similarities, the BPs comprised in the WikiPathways were interactive. Pearson correlation analysis revealed a significant positive correlation among gap junctions, smooth muscle contraction and anti-inflammatory signaling and between acetylcholine binding and downstream events and neurotransmitter release cycle. Finally, VNS exerted its effects on p-pC-MG by reducing the enrichment of human orthologous risk genes for intestinal and extra-intestinal diseases and on p-pC-ISG by decreasing the activation of angiotensin pathway, respectively (Fig. 7b).

**Fig. 7.**
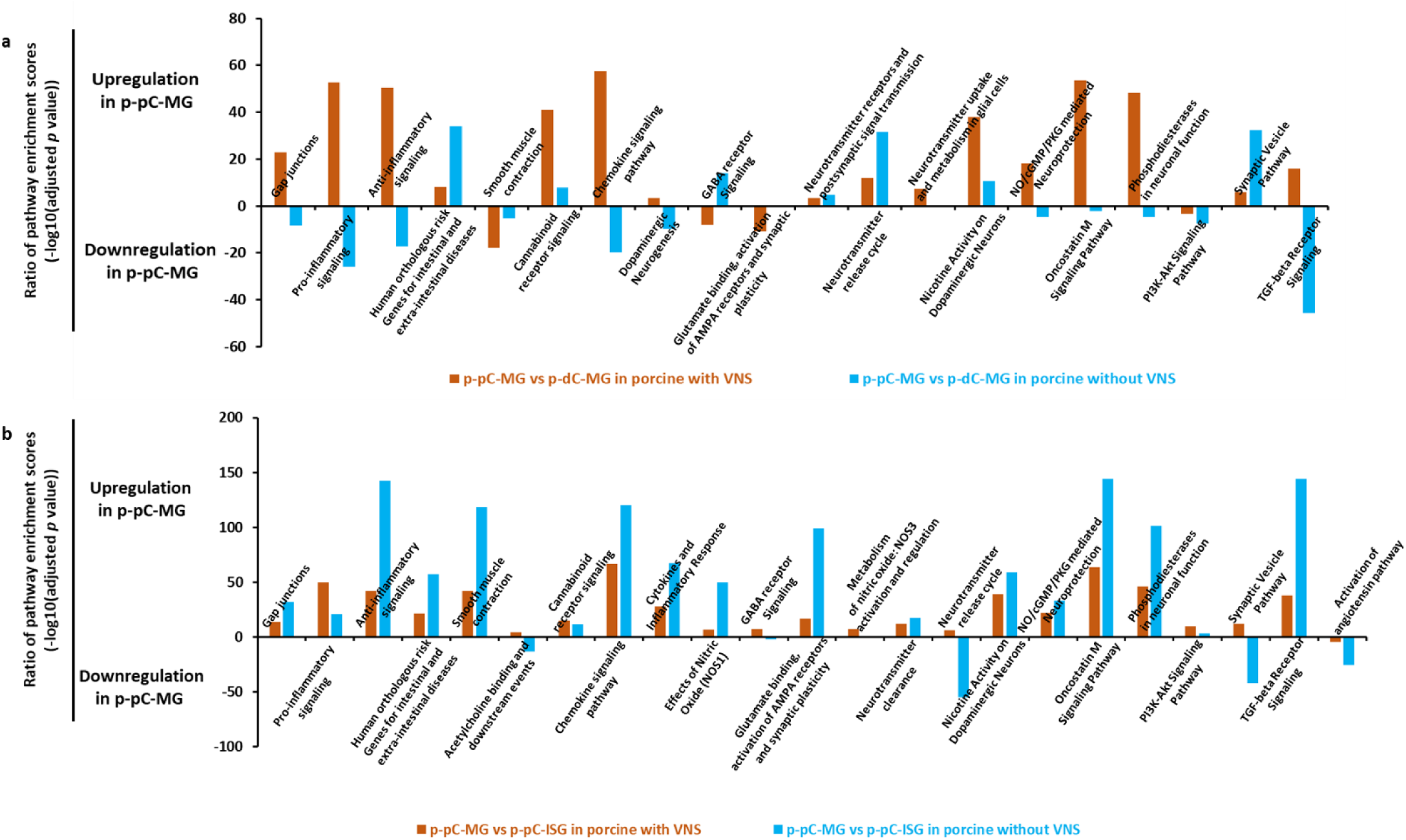
Comparison of pathway enrichment in myenteric ganglia (MG) between porcine proximal and distal colon (p-pC, p-dC) (a), and between MG and inner submucosal ganglia (ISG) in p-pC (b) with and without vagal nerve stimulation (VNS). The figures show the enrichment ratio of the same WikiPathway involving upregulated and downregulated genes in each list of differentially expressed genes based on the bulk RNA sequencing data. The light blue color represents the enrichment ratio without VNS and the orange color represents the enrichment ratio with VNS.

Cell-type contribution to the enrichment of the WikiPathways under VNS was further evaluated based on the comparison of p-pC-MG and p-dC-MG (Fig. 8). The functional linkages involving all cell-type specific DEG that mediated the top five WikiPathways covered >96% of those involving all DEG in each comparison with or without VNS (Supplementary Table 2). In cholinergic neurons of p-pC-MG, VNS substantially increased the enrichment of all top five WikiPathways including chemokine signaling pathway, oncostatin M signaling pathway, TGF-beta receptor signaling, gap junctions and NO/cGMP/PKG mediated neuroprotection. By contrast, VNS reduced the enrichment of gap junctions in SLC41A1_Glia, while increasing those of chemokine signaling pathway in SPC24_Glia. In addition, VNS increased the enrichment scores of oncostatin M signaling pathway in both glutamatergic and nitrergic neurons in p-pC-MG. VNS exerted its effects on glutamatergic neurons in p-dC-MG by increasing enrichment of TGF-beta receptor signaling and decreasing enrichment of chemokine signaling pathway. The enrichment ratio of the shared BPs between the investigated WikiPathways and pro- or anti-inflammatory signaling was calculated to assess the contribution of the pathway enrichment alterations to inflammatory status under VNS. Under VNS, cholinergic and glutamatergic neurons and SLC41A1_Glia contributed to anti-inflammation, while only nitrergic neurons contributed to inflammation in p-pC-MG. By contrast, anti-inflammation was attributed to glutamatergic and nitrergic neurons and SLC41A1_Glia in p-dC-MG (Fig. 9).

**Fig. 8.**
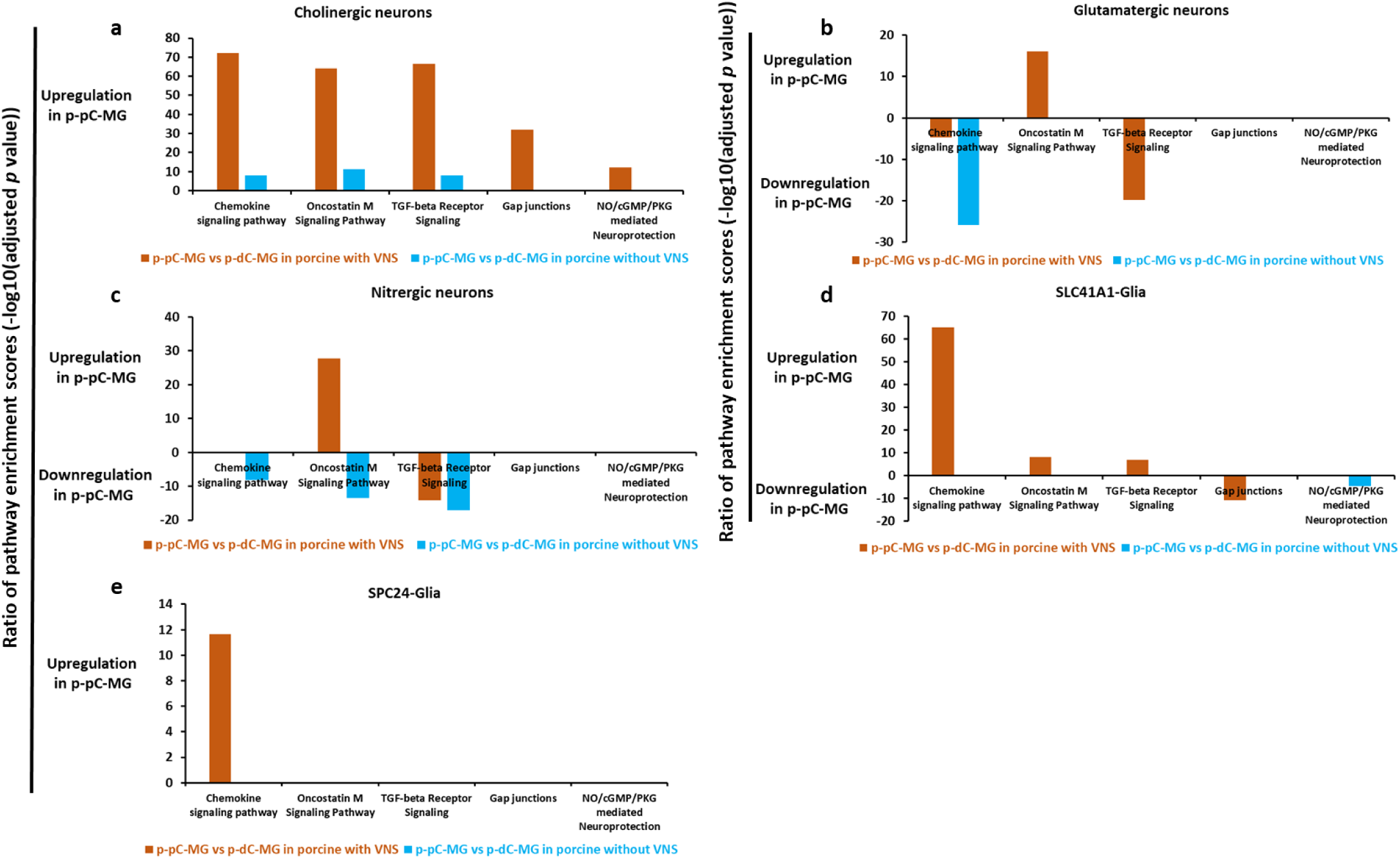
Comparison of cell-type pathway enrichment in myenteric ganglia (MG) of porcine proximal and distal colon (p-pC, p-dC) with and without vagal nerve stimulation (VNS). The figures show the enrichment ratio of the same WikiPathway involving upregulated and downregulated genes in each list of differentially expressed genes and only show the enrichment ratio of the top five WikiPathways. The light blue color represents the cell-type enrichment ratio without VNS and the orange color represents the cell-type enrichment ratio with VNS.

**Fig. 9.**
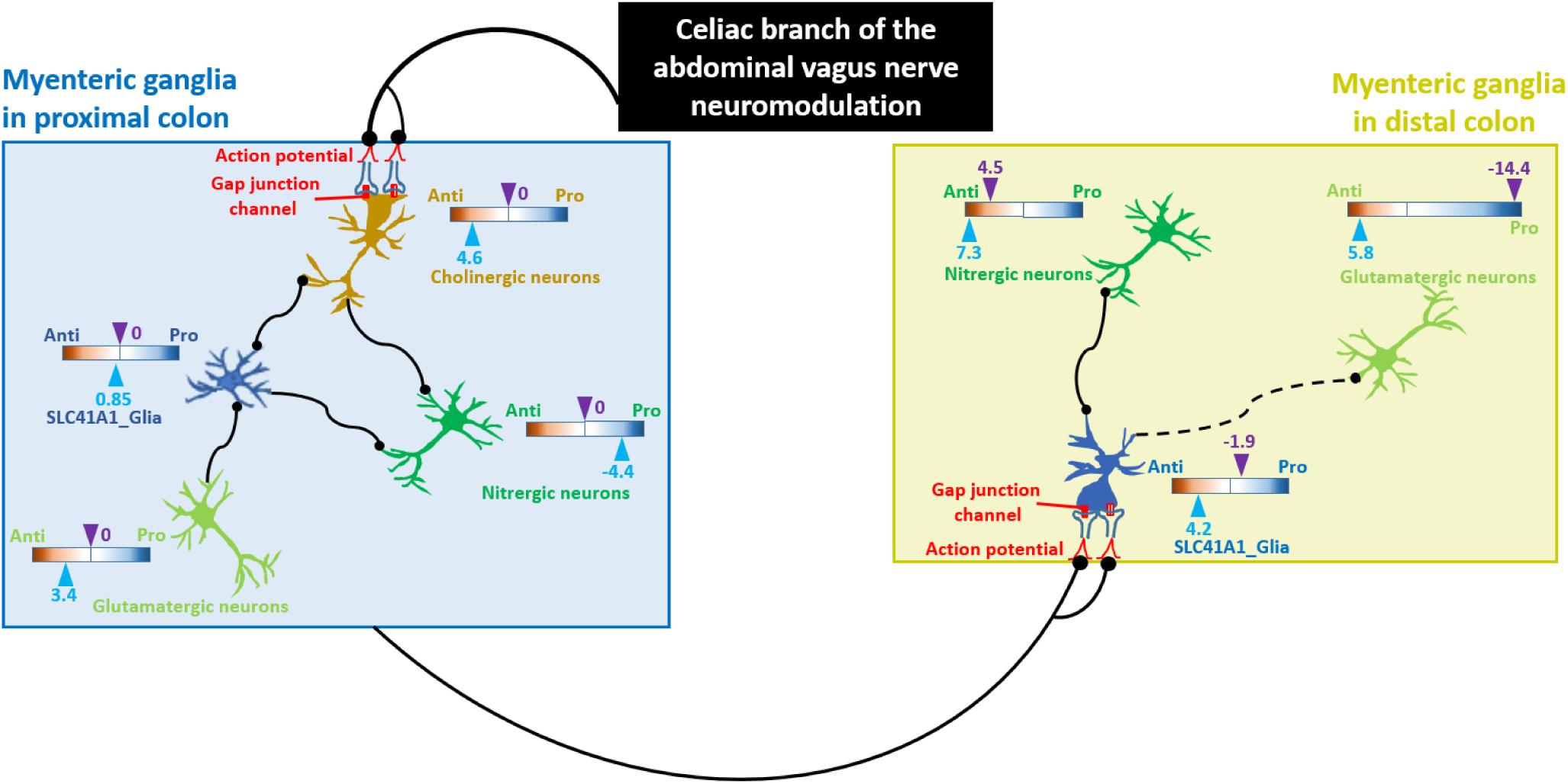
Cartoon diagram illustrating potential electrical transmission along the proximal-distal axis of the porcine colon under vagal nerve stimulation (VNS) and the response mechanisms of certain neuronal/glial cell populations to VNS. The enrichment ratio of the shared biological processes (BPs) between the top five WikiPathways and pro- or anti-inflammatory signaling (pro/anti) was calculated. The solid and dot lines represent cell-cell interactions with and without alterations under VNS, respectively. Black dots represent the targeting sites. Color represents a log10(pro/anti) distribution from Anti (orange) to Pro (blue). Arrowheads in light blue and purple indicate the enrichment ratio with and without VNS, respectively. Pro, pro-inflammation. Anti, anti-inflammation.

The response of cell-cell interactions to VNS was estimated through differential expression of genes encoding ligands and receptors in neuronal and glial subsets (Figs. 9, 10). VNS potentially improved the cell-cell interactions in p-pC-MG, including the interactions between SLC41A1_glia and cholinergic or glutamatergic or nitrergic neurons, cholinergic neurons and nitrergic neurons, and SPC24_glia and cholinergic or nitrergic neurons. The interactions between SLC41A1_glia and cholinergic neurons may be stronger because of the upregulated expression of the genes encoding both ligands and receptors. VNS impacts on p-dC-MG by potentially improving the interactions between cholinergic and glutamatergic neurons and between SLC41A1_glia and nitrergic neurons (Fig. 9, 10).

**Fig. 10.**
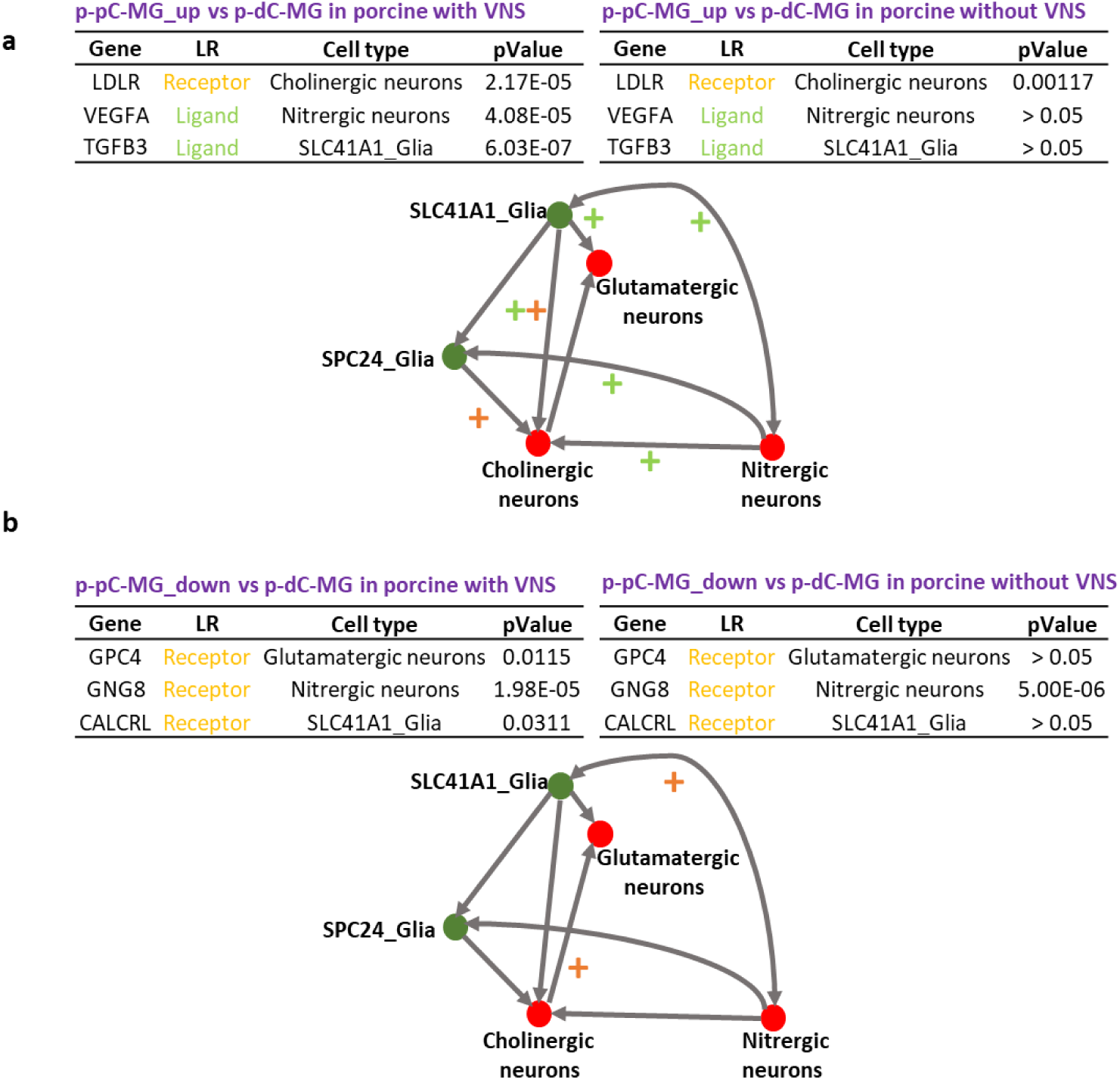
Potential alterations of cell-cell interactions under vagal nerve stimulation (VNS) in comparison of myenteric ganglia (MG) of porcine proximal and distal colon (p-pC, p-dC). The alterations were estimated through differential expression (p-value) of ligands (light green) and receptors (orange) expressed in neuronal and glial subsets in MG with VNS compared with those without VNS. + labels the upregulated interaction of cell subsets where there are the genes expressing ligands/receptors with smaller p-value under VNS.

## Discussion

The cross-comparison of the regional transcriptomic profiling between porcine and human indicates that the myenteric ganglia in proximal or distal colon express a conserved core gene program of orthologous genes from porcine to human with regional heterogeneity. Interestingly, VNS on the porcine modulates the conserved transcriptional program in the myenteric ganglia of proximal or distal colon, which may be implicated in the VNS-altered colonic physiological functions as we reported before^17^, suggesting that human colonic functions and neuromodulation could be modeled in the porcine.

Recent studies have identified several conserved marker genes in ENS cell subtypes between human and mouse based on orthologous gene expression^18,28^. According to our bulk RNA-seq data from both porcine and human ENS, we uncovered 7246 and 2601 conserved genes between h-aC-MG and p-pC-MG (p-value > 0.001) and between of h-dC-MG and p-dC-MG (p-value > 0.001) respectively from a total of 12291 high-quality orthologous genes using R package SCBN. The functional linkages of gene regulatory networks, where single genes are positioned, have been used as powerful indicator, reflecting mechanistic insights and functional similarities across species^29–32^. Pathway analysis could identify biological pathways that are enriched in DEG lists and highlight the overlaps among pathways based on the shared genes^29^. We extracted the gene networks to indicate the differentiation between h-aC-MG and h-dC-MG and between p-pC-MG and p-dC-MG. Twelve porcine orthologous genes from comparison of h-aC-MG and h-dC-MG were found, of which only 2 genes were the conserved genes based on the outputs from R package SCBN. Nonetheless, the 12 orthologous genes served as driver genes, whose networks represented all the characteristics of p-pC-MG and p-dC-MG in porcine, in terms of 100% linkage coverage (Fig. 1). It was worth noting that the networks derived from the 12 porcine orthologous driver genes interpreted >90% functional alterations in p-pC-MG and p-dC-MG of porcine subjected to VNS (Fig. 1). The networks mediated by the human orthologous genes from comparison of p-pC-MG and p-dC-MG did not realize 100% coverage of the linkages derived from the DEG lists in comparison of h-aC-MG and h-dC-MG, probably because of the lack of complete annotation and assembly of the porcine genome^33^. Thus, although there were few conserved genes between porcine and human colon, the networks of interacting orthologous genes orchestrated almost the same complex colonic functions across species, and p-pC and p-dC could serve as predictors, with important implications in understanding of human colonic functions and neuromodulation.

Our work shows for the first time the ENS transcriptomic heterogeneity between MG and ISG and along the proximal-distal axis of the porcine colon. Comparisons of MG and ISG in each of p-pC and p-dC uncovered several pathways responsible for normal colonic physiological functions such as peristalsis and secretion. The inflammatory signaling pathways prevailed in MG, including interactions between immune cells and neurons (e.g., oncostatin M signaling pathway and cytokines and inflammatory response), cytokine production (e.g., chemokine signaling pathway) and anti-inflammation (e.g., TGF-beta receptor signaling) (Fig. 2). Of them, the enrichment of anti-inflammation, involving >99% of the DEG, was the highest. The reason for the inflammatory signaling pathways predominant in MG is probably because the inflammatory bowel disease (IBD) risk genes (e.g., *SMAD3* and *OSMR*) were significantly expressed in MG. Our data also showed that all inflammatory signaling pathways closely interacted with smooth muscle contraction, indicating that the inflammatory status influenced smooth muscle activity. In addition, effects of nitric oxide mediated by *nNOS* and neurotransmitter clearance mediated by *SLC6A4* were dominant in MG. Both nNOS and SLC6A4 that lowers the levels of serotonin are responsible for relaxation of smooth muscle^34–36^. Furthermore, the enrichment of glutamate binding, the activation of AMPA receptors and synaptic plasticity was much higher in MG than in ISG and directly interacted with the BPs such as the regulation of blood circulation and smooth muscle contraction in MG. By contrast, ISG had higher enrichment of activation of angiotensin pathway, synaptic vesicle pathway, acetylcholine synthesis, binding and downstream events and dopamine transport and secretion (Fig. 2), all of which interacted with each other. Angiotensinogen (AGT) that was located in the core of activation of angiotensin pathway and selected in our study has been involved in the development of GI mucosal injury due to SARS-CoV-2 and other factors^37,38^. Therefore, we used the enrichment of this pathway to evaluate the mucosal integrity which showed that only ISG contributed to the modulation of mucosal integrity. Moreover, we found that the activation of angiotensin pathway, acetylcholine synthesis, binding and downstream events and dopamine transport and secretion shared the BPs namely signaling and cell communication, suggesting the potential roles of acetylcholine-dopamine balance in maintenance of mucosal integrity. It is worth noting that neurotransmitter uptake and metabolism in glial cells, mediated by SLC1A3, a glial high affinity glutamate transporter, was dominant in MG only in p-dC (Fig. 2). A steep oxygen gradient along the proximal-distal axis of colon exists, which leads to formation of reactive oxygen species and the resulting glutamate excitotoxicity^39–41^. Glutamate uptake in glia guarantees the neural circuit integrity because SLC1A3 connected the BPs such as synaptic signaling and neurogenesis.

When comparing MG between p-pC and p-dC, we found that the inflammatory signaling pathways such as cytokines and inflammatory response, chemokine signaling pathway and TGF-beta receptor signaling were prevalent in p-dC-MG (Fig. 3). There is evidence that hypoxia and inflammation are intertwined^41^. In addition, p-dC-MG compared to p-pC had much higher enrichment of smooth muscle contraction, NO/cGMP/PKG mediated neuroprotection and dopaminergic neurogenesis (Fig. 3), with all of them having interactions. It has been reported that GUCY1A1 (guanylate cyclases), which was involved in NO/cGMP/PKG mediated neuroprotection, is activated by nitric oxide and plays a key role in inhibiting neuroinflammation^42^. From this perspective, we infer that the combination of nitrergic and dopaminergic neurons contributes to the regulation of smooth muscle contraction. This was confirmed by our scRNA-seq data showing that the anti-inflammatory status, determined by more BPs shared by TGF-beta signaling, chemokine signaling pathway and anti-inflammatory signaling, was closely related to the differentiation of two neuronal populations (Fig. 6). This is consistent with recent reports, implicating the role of TGF-beta in differentiation of nitrergic and catecholaminergic enteric neurons^43,44^. Recent study showed that the interactions of these two neuronal populations could stabilize asynchronous contractile activity and lead to the generation of colonic peristalsis in proximal colon^45^. However, our results suggested that the contribution of dopaminergic and nitrergic neurons to colonic peristalsis is much greater in p-dC than in p-pC. By contrast, p-pC-MG had more complex neuronal wiring, highlighted by the high enrichment of nicotine activity on dopaminergic neurons (mediated by *TH*), acetylcholine synthesis (mediated by *ChAT*), synaptic vesicle pathway, neurotransmitter receptors and postsynaptic signal transmission (mediated by *HTR3A*), neurotransmitter release cycle and GABA receptor signaling (mediated by *GABRB3* and *GABRA4*) (Fig. 3). Cholinergic neurons played leading roles based on their contribution to the top five WikiPathways (Fig. 6), which is in accordance with the findings of Li et al^15^.

The comparison of ISG between p-pC and p-dC revealed that the inflammatory signaling pathways such as chemokine signaling pathway and TGF-beta receptor signaling prevailed in p-dC-ISG, while in p-pC-ISG, there was higher synaptic activity, characterized by the high enrichment of nicotine activity on dopaminergic neurons (mediated by *TH*) and neurotransmitter receptors and postsynaptic signal transmission (mediated by *HTR3A* and *HTR3B*) (Fig. 3). However, we also found high enrichment of blood vessel development and effects of nitric oxide (mediated by *nNOS*) in p-dC-ISG. Nerve fibers may contribute to the perivascular circulation based on the distribution of nNOS and its catalytic activity with nNOS-derived NO maintaining vasodilation^46^. Unlike the comparison of MG between p-pC and p-dC, dopaminergic neurogenesis and dopamine biosynthetic process were predominant in p-pC-ISG. Evidence showed that dopamine could promote colonic mucus secretion^47^.

Our previous study has indicated that the functional sphere of the influence of VNS on porcine colon was pan colonic, from p-pC to p-dC, and VNS increased contractions across the colon^17^. The evidence also showed that the vagal nerve endings often synapse onto neurons in the myenteric plexus^48^. Consistent with this report, our data showed that the major changes induced by VNS happened in MG, especially in p-pC-MG. Although VNS led to some differences of pathway enrichment patterns when comparing p-pC-MG with p-pC-ISG, the alterations were not as prominent as those in comparison to p-pC-MG and p-dC-MG characterized by the enrichment pattern reversal of more than half the investigated pathways (Fig. 7). Compared with p-pC-MG, the obvious changes in p-dC-MG were the enhanced enrichment of smooth muscle contraction and glutamate binding, activation of AMPA receptors and synaptic plasticity, which were interactive. This suggested the crucial roles of synaptic plasticity mediated by glutamate in regulation of smooth muscle contraction. Interestingly, cholinergic neurons contributed to the enrichment of glutamate binding, activation of AMPA receptors and synaptic plasticity, implying that cholinergic neurons may co-release both acetylcholine and glutamate, and the released glutamate could activate postsynaptic neurons (glutamatergic neurons)^49^. Our cell-cell interaction data also confirmed the potentially upregulated connection between cholinergic and glutamatergic neurons under VNS (Fig. 10b). Together with the improved enrichment of GABA receptor signaling, we speculated that VNS promoted the occurrence of rhythmic phasic contractions and fecal pellet expulsion in p-dC. However, in p-pC-ISG, there was the decreased enrichment of acetylcholine binding and downstream events, neurotransmitter release cycle and synaptic vesicle pathway. Of them, VNS led to the highest alterations in the enrichment of glutamate release cycle (*GLS*/*GLS2*). This may be related to the VNS-induced the activation of an anti-inflammatory response^40,41^. This is in accord with the improvement of the mucosal integrity, characterized by the reduced enrichment of activation of angiotensin pathway (Fig. 7b).

Gap junctions greatly contributes to electrical transmission^50^. Here, we used the enrichment changes of gap junctions to interpret the influences of VNS on the transcriptomic profiling. When comparing p-pC-MG with p-pC-ISG, the enrichment of gap junctions, anti-inflammatory signaling and smooth muscle contraction was surprisingly reduced in p-pC-MG by VNS (Fig. 7b) and had a significant positive correlation (Supplementary Fig. 5). In addition, we found that glutamate release cycle and gap junctions showed the highest similarity score, suggesting that glutamate transmission may complement the function of gap junctions. Consequently, glutamate release triggered acetylcholine binding and downstream events (*CHRNA5*) due to their significantly positive correlation (Supplementary Fig. 5). The potentially synergistic functions of glutamate and acetylcholine may be important for functional modulation in p-pC-MG. Collectively, the alterations of the pathway enrichment under VNS may reduce disease risks in view of the decreased enrichment of human orthologous risk genes for intestinal and extra-intestinal diseases, associated with gut dysmotility and inflammation in p-pC-MG (Fig. 7b).

Compared with p-dC-MG, enrichment of gap junctions was highly increased in p-pC-MG, while enrichment of synaptic vesicle pathway, neurotransmitter receptors and postsynaptic signal transmission and neurotransmitter release cycle was decreased. We speculated that there is direct mechanistic relationship between the formations of neuronal gap junction coupling and the disappearance of chemical transmission in most neuronal cell types as reported^50^. Previous studies showed that VNS reduces intestinal inflammation through the cholinergic anti-inflammatory pathway^51^. In line with the statement, we found that the enrichment ratio of pro- to anti-inflammatory signaling (pro/anti) in porcine without VNS was 1.5 times higher than that in porcine with VNS, suggesting the anti-inflammatory effects of VNS (Fig. 7a). The enrichment of pro-inflammatory signaling under VNS was attributed to some extent to nitrergic neurons as supported by a recent report showing that NOS1 led to an outburst of inflammatory reactions in macrophages via activator protein-1^52^. Whereas, enrichment of anti-inflammatory signaling was attributed to both cholinergic and glutamatergic neurons and SLC41A1_Glia in p-pC-MG (Fig. 8, 9). The enteric glia were thought to be essential to reduce the intestinal inflammatory response under VNS^53^. In the vagal circuitry, glutamate is a primary neurotransmitter involved in key gastrointestinal functions^54^. However, its anti-inflammatory effects are rarely reported except that excessive release of glutamate into the extrasynaptic space led to inflammatory responses, which could be corrected by glial cells in CNS^55^. Our result indicated that the synergies of glutamatergic neurons with enteric glia were also potentially present in ENS under VNS because of the increased enrichment of neurotransmitter uptake and metabolism in glial cells (GLUL, conversion of glutamate and ammonia to glutamine) p-pC-MG (Fig. 7a). Duo to the direct interactions between smooth muscle contraction and anti-inflammatory signaling, the anti-inflammatory properties indeed reduced the enrichment of human orthologous risk genes for intestinal and extra-intestinal diseases, associated with gut dysmotility and inflammation (Fig. 7a). We found that *OSMR*, the only risk gene for IBD was expressed significantly in p-pC-MG compared to p-dC-MG under VNS. *OSMR* was enriched in SLC41A1-Glia and the *OSMR*-mediated pathways shared BPs with pro- and anti-inflammatory signaling. The enrichment ratio of the shared BPs (pro/anti) was 1.3 in porcine with VNS, while pro/anti was 1.2 in porcine without VNS, which did not show the significant anti-inflammatory effects of VNS on IBD. Future work should involve more risk genes for IBD to convincingly identify the exact correlation between vagal function and IBD. Finally, we derived the VNS influences on p-dC-MG from the improved enrichment of gap junctions in SLC41A1-Glia of p-dC-MG, compared with p-pC-MG (Figs. 8d, 9). Surprisingly, unlike the pro-inflammatory impacts of nitrergic neurons in p-pC-MG, they played the anti-inflammatory roles in p-dC-MG, together with glutamatergic neurons and SLC41A1_Glia. Moreover, the contribution of SLC41A1_Glia to anti-inflammation was much higher in p-dC-MG. Interestingly, the effects of VNS on glutamatergic neurons and SLC41A1_Glia in p-dC-MG led to switch of their contribution from inflammation to anti-inflammation (Fig. 9). Different from the attribution of glutamatergic neurons to anti-inflammation in p-pC-MG under VNS, the increased enrichment of glutamate binding, activation of AMPA receptors and synaptic plasticity in cholinergic neurons ensured the correction of glutamatergic neurotransmission dysfunction in p-dC-MG (Fig. 7a). Thus, we inferred that the synergies of glutamatergic neurons with cholinergic neurons contributed to anti-inflammation in p-dC-MG.

In summary, this study compared for the first time the transcriptomic profiles in the colonic ENS between porcine and human and across two layers of enteric plexus along the proximal to distal colonic regions in porcine. The regional-specific gene programs were identified at both bulk and single-cell resolution in porcine and regional-dependent highly conserved core transcriptional programs were revealed between porcine and human. VNS exerted its effects mainly on the myenteric ganglia in porcine. Regional-specific transcriptomic responses to VNS shared >90% of the conserved core transcriptional programs between porcine and human. Our study provides a fundamental resource for better understanding of the importance of porcine model in the colonic translational research.

## Methods

### Tissue collections

#### Porcine colon

Six castrated male and three intact female Yucatan minipigs (∼7 months old, 25-36 kg, S&S Farms, Ramona, CA) were fasted for 12 h and anaesthetized by intramuscular application of midazolam (1 mg/kg, cat # 067595, Covetrus, Dublin, OH), ketamine (15 mg/kg, cat # 068317, Covetrus, Dublin, OH) and meloxicam (0.3 mg/kg, #049755, Covetrus, Dublin, OH). Three out of the six male animals underwent electrical stimulation of the celiac branch of the abdominal vagus nerve (2 Hz, 0.3-4 ms, 5 mA, 10 min) using pulse train. A detailed experimental protocol for VNS is available at https://www.protocols.io/view/tache-mulugeta-ot2od024899-colon-tissue-electrical-3rmgm. The colonic specimens with full thickness (about 1.5 cm long) were removed from the proximal colon (p-pC, about 10 cm from the ceco-colic junction), transverse colon (p-tC, about 10 cm from the end of the proximal, specifically about 10 cm from the end of the centrifugal spiral colon) and distal colon (p-dC, about 20 cm from the ano-rectum). Time intervals between anesthesia injection and tissue collections were 20-30 min for naïve pigs without VNS and about 5 h for these with VNS. All samples were embedded in OCT, snap-frozen in dry ice and stored at -80°C for LCM procedures. One extra piece (4×4 cm) of full thickness colon tissue samples was harvested from each p-pC, p-tC and p-dC of 4 naïve porcine (2 of each sex) and processed for scRNA-seq. All animal care and procedures were performed following National Institutes of Health guidelines for the humane use of animals, with approval and in accordance with the guidelines of the University of California at Los Angeles (UCLA) Institutional Animal Care and Use Committee, Chancellor’s Animal Research Committee (ARC) (protocol 2018-074-01). All efforts were made to avoid suffering.

#### Human colon

The full thickness of colonic specimens (about 5×3 cm) were dissected from healthy margin of the post-operative ascending, transverse and descending colon (h-aC, h-tC, h-dC, 4 of each) from 12 patients (4 males and 8 females, median age 47 years old, range 35-66 years) with colonic adenocarcinoma. All specimens were provided by the UCLA Translational Pathology Core Laboratory after dissection and examination to be normal under macro- and microscopic inspection. None of the patients had active colonic infections when the tissues were collected. Samples were immersed in Belzer UW^®^ Cold Storage Solution (Bridge to Life Ltd, Columbia, SC) on ice. Once delivery, samples were instantly embedded in OCT, snap-frozen in dry ice and stored at -80°C for LCM procedures. The interval time between colonic resection and snap freezing was 70-90 min. The use of human colon tissues was approved by the UCLA Institutional Review Board for Biosafety and Ethics (IRB #17-001686). Informed consent has been obtained in all cases.

### Laser-capture microdissection (LCM) of enteric ganglia from human and porcine colon and bulk RNA-seq

A range of 25-40 ganglia/subject were harvested from ISG and MG respectively of p-pC, p-tC and p-dC (6 without VNS and 3 with VSN) and from MG of h-aC, h-tC and h-dC (12 subjects) using LMD-6000 Laser Micro-dissection System (Leica Microsystems, Wetzlar, Germany) (Supplementary method 1). Total RNA was extracted for construction of cDNA libraries (Supplementary method 2) and sequenced on an Illumina HiSeq 3000 sequencer as 50 base pair single-end reads. All of the reads were then aligned to the sus_scrofa or human genome using STAR and the DEG lists were generated using edgeR (Supplementary method 3) and used for pathway enrichment analysis.

### scRNA-seq in colonic enteric ganglia of naïve porcine

The mucosa and submucosa were peeled off from each p-pC, p-tC and p-dC of 4 naïve porcine (2 of each sex). The muscularis externa containing myenteric ganglia were processed for cell suspension preparation (Supplementary method 4). ∼10,000 single cells per cell suspension were loaded on the 10x Genomics Chromium platform and sequenced on an Illumina NextSeq 500 Sequencer (75 cycles). Sequencing was performed in paired-end mode. Cell Ranger 3.1.0 count function was used to align and quantify the reads against sus_scrofa genome assembly, created using the Cell Ranger 3.1.0 mkref function.

### Identification of cell types from scRNA-seq data and analysis of ligand-receptor interaction in colonic enteric ganglia of naïve porcine

The scRNA-seq data were processed for cell selection, filtration, normalization, scaling, cell cluster identification and annotation of neuronal and glial clusters using R toolkit Seurat (Supplementary method 5). The neuronal and glial populations acquired from 12 scRNA-seq datasets were used to further gain the subpopulations of neurons and glia using Seurat. A total of 511, 286 and 443 neurons and 316, 318 and 389 glia were isolated from p-pC, p-tC and p-dC respectively, from which 3 neuronal (cholinergic, nitrergic and glutamatergic) and 2 glial subpopulations (marked by *SLC41A1* and *SPC24*, respectively) were identified. Cell-cell interactions were inferred from the single-cell transcriptomic data based on the ligand-receptor (L-R)-pair list in porcine generated using R package ‘LRBase.Ssc.eg.db’. Firstly, scRNA-seq data were read using R package ‘DropletUtils’ and then the target cells were selected using unique molecular identifiers of neurons and glia. After data normalization, the R package ‘scTGIF’ was used to convert Ensembl ID to NCBI gene ID and a SingleCellExperiment object was created using R package ‘SingleCellExperiment’. At last, the R package ‘scTensor’ was used to detect and visualize cell-cell interactions.

### Single-molecule fluorescence in situ hybridization (smFISH)

To validate the key discriminatory markers for the subsets of neuronal and glial cells clustered from scRNA-seq data, we performed RNAscope, a smFISH technique with fresh-frozen sections (12 µm) crossing the myenteric ganglia in p-pC. RNAScope Multiplex Fluorescent Kit v2 (Advanced Cell Diagnostics) was used per manufacturer’s recommendations. The hybridization detection was carried out using Opal Fluorophore Reagent Packs Opal-520 (channel (C) 1)/570 (C2)/690 (C3) (1:1500) (Akoya Biosciences). Probes (Advanced Cell Diagnostics) used for smFISH include *GAP43* (1076491-C1) for annotating enteric neurons validated with *ELAVL4* (1076501-C2) encoding Hu C/D, a well-known pan neuronal marker; *CLDN11* (1076511-C1) for annotating enteric glial cells, *SLC41A1* (1076531-C1) and *SPC24* (1076541-C2) for annotating two subtypes of enteric glial cells, validated with *GFAP* (1039661-C3) encoding glial fibrillary acidic protein (GFAP), a well-known marker for pan enteric glial cells; *ACLY* (1076551-C2) for annotating cholinergic neurons validated with *ChAT* (849981-C1) encoding choline acetyltransferase (ChAT), a well-known marker for cholinergic neurons; *GLS* (1076571-C1) for annotating glutamatergic neurons validated with *SLC17A6* (1076581-C2) encoding vesicular glutamate transporter 2 (VGLU2), a well-known marker for glutamatergic neurons; *ARHGAP18* (107659-C1) for annotating nitrergic neurons validated with *NOS1* (1076601-C2) encoding neuronal nitric oxide synthase (nNOS or NOS1), a well-known marker for nitrergic neurons. A 3-plex RNAscope® positive control including 3 probes for porcine *ACTB* (1076681-C1), *PPIB* (428591-C2) and *GAPDH* (1076691-C3), three housekeeping genes encoding β-actin (ACTB), peptidylprolyl isomerase B (PPIB) and glyceraldehyde 3-phosphate dehydrogenase (GAPDH), and a 3-plex negative control containing 3 probes with C1, C2 and C3 (Advanced Cell Diagnostics) were served to test the specificity of RNAscope assay and the qualities of tissue samples. Images were taken using a Zeiss LSM 710 confocal microscope (Carl Zeiss Microscopy, LLC, White Plains, NY) with a laser set of 488 (C1)/561 (C2)/647 (C3). Five to ten optical sections (Z-stack) crossing the myenteric ganglia with frame 436×436 μm and 1 μm apart (20× objective) were acquired and overlaid using Imaris 9.7 for Neuroscientists (Bitplane Inc., Concord, MA).

### Deconvolution of bulk RNA-seq data using scRNA-seq datasets from colonic enteric ganglia of naïve porcine

The R package ‘SCDC’ was used to deconvolve the bulk RNA-seq data from the myenteric ganglia of naïve porcine colon with an ENSEMBLE method using default settings^24^. The 12 scRNA-seq datasets generated in this study were served as references. With marker genes specific for the specified cluster, five subpopulations of cells including cholinergic, nitrergic, glutamatergic neurons and two types of glia were identified in scRNA-seq datasets and selected to construct basis matrix in p-pC, p-tC and p-dC. For each scRNA-seq dataset with raw counts, a quality control procedure was carried out to filter the cells with the threshold of keeping a single cell from an assigned cluster of 0.7. The grid search method was used to derive the ENSEMBLE weights with the search step size of 0.01. SpearmanY from the grid search result represented the maximum of the Spearman correlation between observed cell-type gene expression in scRNA-seq datasets and predicted cell-type gene expression (probabilities) in bulk RNA-seq data. The cell-type gene lists were then matched to the DEG lists (*p* < 0.05) that were obtained from bulk RNA-seq data analysis. In this way, the cell-type DEG lists were generated and used for pathway enrichment analysis.

### Selection of orthologous genes

A gene ortholog table was first established using human genome as the reference gene list to compare transcription between human and porcine. Gene homology search was performed, using ensemble multiple species comparison tool (http://www.ensembl.org/biomart/). A high-quality orthologous gene list was extracted with whole genome alignment score above 75^56^, resulting in 12291 genes.

### Expression levels and cross-species normalization

Based on the reads of the high-quality ortholog genes, the Python/bioinfokit (v0.9.1) package was used to calculate standard RPKM (reads per kilobase of exon model per million mapped reads) expression values for the orthologous gene set, followed by log_2_ transformation. The cross-species normalization was completed according to the scaling procedure reported by Brawand et al.^57^. Briefly, among the genes with expression values in the inner quartile range, the genes with the most conserved ranks among samples were identified and their median expression levels in each sample were assessed. The scaling factors were derived by adjusting these medians to a common value and used to scale expression values of all genes in the samples. After normalization, empirical cumulative density function (ECDF) (R/ecdf package) was used to estimate their probability distributions, and both Kolmogorov-Smirnov test (Python/scipy.stats.ks_2samp) and Pearson’s Chi-squared test (Python/bioinfokit v0.9.5) were used to evaluate their goodness of fit. R/cor package was applied to compute the cross-species Spearman’s Correlation, which was visualized using R/corrplot package.

### Analysis of pathway enrichment using bulk and single-cell RNA sequencing data

The DEG lists were generated based on the transcriptomic comparisons in both human and naïve porcine samples with datasets pooled from males and females. Those from the male porcine were only used to assess the effects of VNS on the transcriptomic profiling in porcine colon. In order to remove the potential contamination from intestinal muscle during LCM, unique smooth muscle marker genes derived from scRNA-seq were deleted from each DEG list based on bulk RNA-seq data. The modified DEG lists were imported into ClueGO v.2.5.6 in Cytoscape v.3.8.2 with an FDR (false discovery rate) *p*-value cut-off of 0.05 to create and visualize a functionally grouped network of biological processes. In addition, WikiPathways were chosen to provide intuitive views of the multiple interactions underlying biological processes. We searched WikiPathways of interest from human database (https://www.wikipathways.org/) to interpret the data from the porcine colon. The porcine DEG lists were matched to the generated high-quality ortholog gene list. Those involved in the WikiPathways of interest were used to perform enrichment analysis. WikiPathways of interest were classified into 9 categories (see details in Supplementary Table 3).

Some WikiPathways such as smooth muscle contraction and activation of angiotensin pathway were considered to evaluate the influences of MG and ISG on the physiological status of smooth muscle and mucosa. The WikiPathways such as gap junctions associated with electrical stimulation were also included. In addition, pro-/anti-inflammatory signaling (see details in Supplementary Table 4) was used to assess the effects of the vagal electrical stimulation on the inflammatory status. Firstly, the DEG lists were imported into g:Profiler (https://biit.cs.ut.ee/gprofiler/) or ClueGO v.2.5.6, followed by creation and visualization of gene networks in EnrichmentMap v.3.2.1 or ClueGO v.2.5.6 in Cytoscape v.3.8.2 with an FDR *p*-value cut-off of 0.05. The WikiPathways-matching DEG lists were inputted into EnrichmentMap v.3.2.1 or ClueGO v.2.5.6 to compute the FDR *p*-value of the specified WikiPathways using R/harmonicmeanp package. Details can be found in (Supplementary Note 1). The bubble plots were generated using R/ggplot2 package. Similarity matrix was generated in R/rrvgo package at default setting using DEG lists from comparison of p-pC-MG and p-pC-ISG in porcine with VNS. To confirm further whether the findings in porcine have relevance to translational research, the porcine orthologous DEG list from human h-aC-MG vs h-tC-MG, h-aC-MG vs h-dC-MG or h-tC-MG vs h-dC-MG was imported into the EnrichmentMap that was established based on the regionally corresponding DEG list from porcine p-pC-MG vs p-tC-MG, p-pC-MG vs p-dC-MG or p-tC-MG vs p-dC-MG to assess the extent to which each two gene networks are overlapping. In the same way, gene network overlap was assessed based on the human orthologous DEG list from the regional comparisons in porcine and the regionally corresponding DEG list from human or the regionally corresponding DEG lists from porcine with and without VNS.

### Defining the ortholog of human disease risk genes in the porcine colonic ENS

The human disease risk genes relevant to Hirschsprung’s disease, inflammatory bowel disease, autism spectrum disorders, and Parkinson’s disease were selected based on the literatures^58–61^. Their orthologous genes were defined in the porcine colonic ENS and applied for genetic risk prediction of human diseases. Enrichment analysis of WikiPathways that regulate the expression of the disease risk genes was performed based on the above-mentioned procedure.

## Data availability

RNA sequencing data from this study have been deposited into the National Center for Biotechnology Information Gene Expression Omnibus under accession number: GSE197106.

## Acknowledgements

The present work was supported by the NIH SPARC OT2 grant OD024899 (PQY, YT, MM), UCLA/Digestive Diseases Research Center Core P30 DK41301 and Animal Model Core (YT, MM). The authors would like to thank Dr. Weizhe Hong, UCLA Department of Biological Chemistry and Department of Neurobiology for sharing Act-seq protocol and helpful comments. The authors thank UCLA Technology Center for Genomics & Bioinformatics (TCGB) for RNA sequencing service. The authors would also like to acknowledge Dr. Bob Goldberg, UCLA Molecular, Cell, & Developmental Biology for providing LMD6000 facility.

## Author contributions

PQY conceived and designed the project and performed experiments; TL designed, conducted major experiments and analyzed the data; MM (Million Mulugeta) and ML supported the tissue collection and vagal nerve stimulation; MM (Marco Morselli) assisted in cDNA library construction; ST performed alignment for the reads from parts of bulk RNA-seq raw data; TL, PQY wrote the manuscript; YT, MP, MM (Million Mulugeta), MM and ST gave input on the written manuscript; YT supervised the project. All authors listed provide approval for publication.

