## supplementary for "Comparative transcriptomics reveal highly conserved regional programs between porcine and human colonic enteric nervous system"

**Supplementary Methods**

**Supplementary method 1: Laser Capture Microdissection (LCM) and RNA extraction**

Full thickness colon tissue samples were collected from each p-pC, p-tC and p-dC of porcine and from each h-aC, h-tC and h-dC of human. After washed in the ice-cold, DEPC treated PBS, the tissue samples were embedded in OCT, snap-frozen in dry ice and stored at -80°C until cryosectioning. Tissue sections of 10 μm were cut using the Microm HM 500 M cryostat (Micron Instruments, CA, USA) at -20°C and mounted onto PEN membrane (2.0 µm) slides (Leica Microsystems, Wetzlar, Germany) for UV LCM (Leica LMD6000, Leica Microsystems, Wetzlar, Germany). After mounting the sections, the tissue sections were stained with 1% cresyl violet stain solution (Sigma-Aldrich, MO, USA) before undergoing dehydration through graded alcohols (Thermo Fisher Scientific, NY, USA) (75%, 95%, 100%, 100%) and xylene (Sigma-Aldrich, MO, USA) for a total of 3 min and air-dry for 5 min. Tissue sections were microdissected on a UV laser-based Leica LMD6000 laser microdissection system using the following parameters: 20 × (objective), 50 (laser power), 10 (aperture), 8 (speed), 30 (specimen balance). The cutting followed the indicated marks that outlined the desired ISG and MG. We collected 25-40 ganglia per tissue section from ISG and MG in porcine and from MG in human on the 0.5 ml tube cap (USA Scientific, Inc., FL, USA) filled with lysis solution from the QIAgen RNAeasy Micro Kit (Qiagen, CA, USA). Following LCM, total RNA was extracted using the same kit according to the manufacturer’s instructions. All the RNA samples were stored in nuclease-free tubes at -80°C. The quality and quantity of RNA samples were checked using Agilent RNA 6000 Pico Kit (Agilent, CA, USA) on Agilent 2100 Bioanalyzer (Agilent, CA, USA).

**Supplementary method 2: cDNA library construction**

The quality of RNA samples (67 samples) with RIN over 6 were used to construct cDNA libraries using SMART-Seq® Stranded Kit (Takara Bio USA, Inc., CA, USA), whose quality and quantity were checked using Agilent High Sensitivity DNA Kit on Agilent 2100 Bioanalyzer and using Qubit™ dsDNA HS Assay Kit (Thermo Fisher Scientific, NY, USA) on Qubit 2.0 Fluorometer (Thermo Fisher Scientific, NY, USA), respectively, according to manufacturer protocol. SMART-Seq® Stranded Kit is designed to analyze degraded, partially degraded or high-integrity RNA and deplete ribosomal cDNA.

**Supplementary method 3: Read alignment and generation of DEG using bulk RNA-seq data**

The libraries were sequenced on an Illumina HiSeq 3000 sequencer as 50 base pair single-end reads at UCLA Technology Center for Genomics & Bioinformatics (TCGB). Sus_scrofa genome files (.fa) and annotation file (.gtf) were downloaded at https://uswest.ensembl.org/Sus_scrofa/Info/Index, which were used to create the sus_scrofa genome index using STAR v2.7.1a^1^. Human reads were aligned to Ensembl release 97 human with STAR. The fastq files from TCGB were then aligned against the genome assembly using STAR, followed by assessment for the total number of aligned reads and total number of uniquely aligned reads to evaluate sequencing performance. The gene counts were then imported into R package edgeR^2^, a count-based statistical method, and trimmed mean of M-values (TMM) normalization size factors were calculated to adjust for differences in library size across samples. The TMM size factors and the matrix of counts were imported into the R package Limma, and a weighting approach with the voomWithQuality-Weights function and an additive generalized linear model in Limma were used to deal with variations in sample quality and to correct batch effects created by library preparation and sequencing of large numbers of samples over time, respectively^3^.

**Supplementary method 4: Cell suspension preparation from naïve porcine colon**

One extra piece (4×4 cm) of full thickness colon tissue samples was collected from each p-pC, p-tC and p-dC of 4 naïve porcine. The tissue samples were washed in ice-cold, carbogen (95% O_2_ and 5% CO_2_)-bubbled PBS. The muscularis externa containing myenteric ganglia were peeled off from the underlying tissue using forceps, followed by incubation in enteric neuron media, containing Neurobasal A media with B-27 (Thermo Fisher Scientific, NY, USA), 2 mM L-glutamine (Thermo Fisher Scientific, NY, USA), 1% fetal bovine serum (FBS) (Thermo Fisher Scientific, NY, USA), 10 ng/ml Glial Derived Neurotrophic Factor (Cedarlane Corporation, NC, USA) and 1× Antibiotic/Antimycotic (Thermo Fisher Scientific, NY, USA), in the presence of 45 µM Actinomycin D (ActD)^4^ (Thermo Fisher Scientific, NY, USA) for 15 min on ice. The samples were then cut into small pieces < 1 mm and transferred to the enteric neuron media containing 1mg/ml collagenase B (Sigma-Aldrich, MO, USA) and dispase II (Sigma-Aldrich, MO, USA) and 45 µM ActD for 1 hour at 37°C. After the addition of 1 mg/ml deoxyribonuclease I (Sigma-Aldrich, MO, USA) and 10% FBS, the tissue pieces were dissociated using fire polished Pasteur pipettes. Following manual trituration, the cells were filtered through 70 µm Nitex mesh filter (Miltenyi Biotec Inc., CA, USA) and pelleted at 375 g for 10 min, and resuspended in the ice-cold, carbogen-bubbled rinse medium containing F12 media (Thermo Fisher Scientific, NY, USA) with 10% FBS and 1× antibiotic/antimycotic. Importantly, the solutions used in all steps were equilibrated in the carbogen gas. Estimation of viable cell number was done by Trypan Blue dye (Thermo Fisher Scientific, NY, USA) exclusion method. After cell staining with DAPI (Thermo Fisher Scientific, NY, USA), the viable cells were collected via fluorescence-activated cell sorting (FACS) (BD FACSAriaII, BD Biosciences, CA, USA) at UCLA Jonsson Comprehensive Cancer Center, whose viability was assessed using a Countess II FL Automated Cell Counter (Thermo Fisher Scientific, NY, USA) at UCLA TCGB. In order to determine the concentration of ActD, after cell suspension preparation from muscularis externa of porcine proximal colon, more than 1 million cells per sample were collected for bulk RNA-seq on an Illumina HiSeq 3000 sequencer and the same strategy was applied to analyze the data (Supplementary method 3). Immediate-early genes (IEGs) are rapidly and transiently induced by various cellular stimuli, which were chosen based on Wu et al^4^. The expression levels of total 127 IEGs were extracted (See Supplementary Fig. 6). The heatmap was generated based on the average gene expression levels using R package ‘heatmap.2’.

**Supplementary method 5: Identification of neuronal and glial clusters from scRNA-seq data using Seurat**

After the selection and filtration of cells (genes expressed in at least 3 cells, cells with reads quantified for between 200 and 2500 genes and percentage of counts coming from mitochondrial genes less than or equal to 5%), data normalization and scaling were performed using default options. Data were batch-corrected using Seurat’s integration method. A K-nearest neighbor approach was employed to identify clusters using the top 10 principal components of the processed expression data with resolution set at 0.5. The UMAP algorithm was used for dimensionality reduction. Marker genes were identified by determining the average log-fold change of expression of each cluster compared to the rest of the cells using Seurat’s FindAllMarkers function using the default settings for the Wilcoxon rank sum test. We identified marker genes as those with an average log-fold change above 0.25. Clusters were labeled using cell types associated with the identified marker genes.

**Supplementary Notes**

**Supplementary Note 1: Analysis of pathway enrichment using bulk and single-cell RNA sequencing data**

Enrichment analysis for biological processes (BPs) was performed using g:Profiler (<https://biit.cs.ut.ee/gprofiler/>) or ClueGO v.2.5.6 according to the protocol presented by Reimand et al.^5^ and Bindea et al.^6^. ClueGO provides predefined selection criteria of representative pathways. For GO levels: 1-4, minimum 50 genes/term were defined and minimum 3 genes/term and 1 gene/term were defined for GO levels: 3-8 and for GO levels: 7-15, respectively. For each DEG list, we selected the minimum GO level, where there were at least 3 BPs at a FDR *p*-value cut-off of 0.05. g:Profiler found the genes that were significantly enriched in BPs using a Fisher’s exact test and multiple-test correction. The results from g:Profiler or ClueGO v.2.5.6 were inputted into EnrichmentMap v.3.2.1 or ClueGO v.2.5.6 in Cytoscape v.3.8.2^7^ to visualize the networks of BPs. All annotations of enriched BPs with a FDR *p*-value cut-off of 0.05. We removed some annotations of BPs that were not correlative with ENS functions such as terms related to heart function. Edge width represented the correlation coefficient between nodes and the similarity cutoff overlap coefficient was set to 0.25.

**Supplementary Figure 1:**

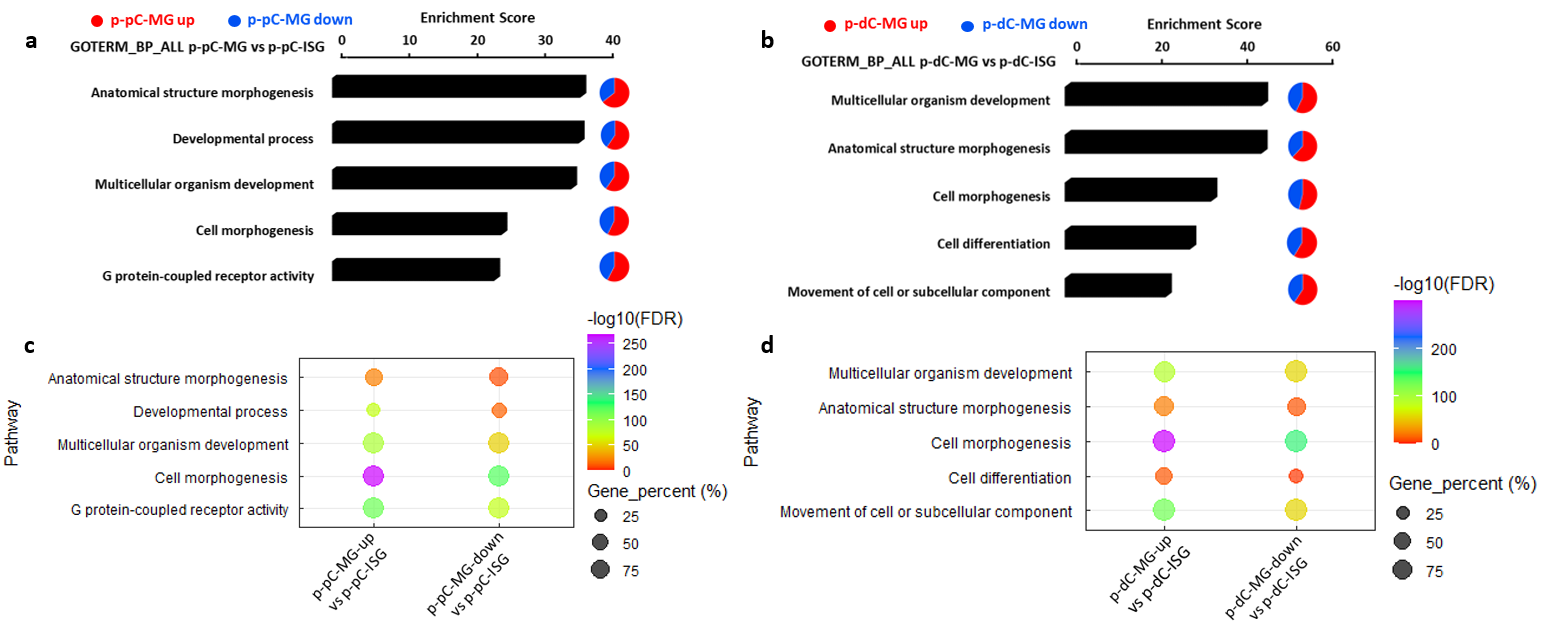

**Supplementary Fig. 1** Comparison of pathway enrichment between myenteric ganglia (MG) and inner submucosal ganglia (ISG) in porcine proximal or distal colon (p-pC, p-dC). a, b Enrichment of top five biological processes (BPs). The pie charts show the percentage of visible genes involved in the specified BPs. Red and blue color represents the genes with upregulated and downregulated expression levels in MG in comparison with ISG, respectively. X-axis marks enrichment score, with the significance cut-off marked by the vertical white line (*p*-value < 0.05). c, d The bubble plots show the gene percentage and enrichment of the top BPs. Circle size represents the ratio of the number of pathway-specific differentially expressed genes (DEG) and the number of total DEG in each DEG list (Gene_percent). Color represents a -log10(FDR) distribution from big (orange) to small (purple). Up, upregulated expression. Down, downregulated expression.

**Supplementary Figure 2:**

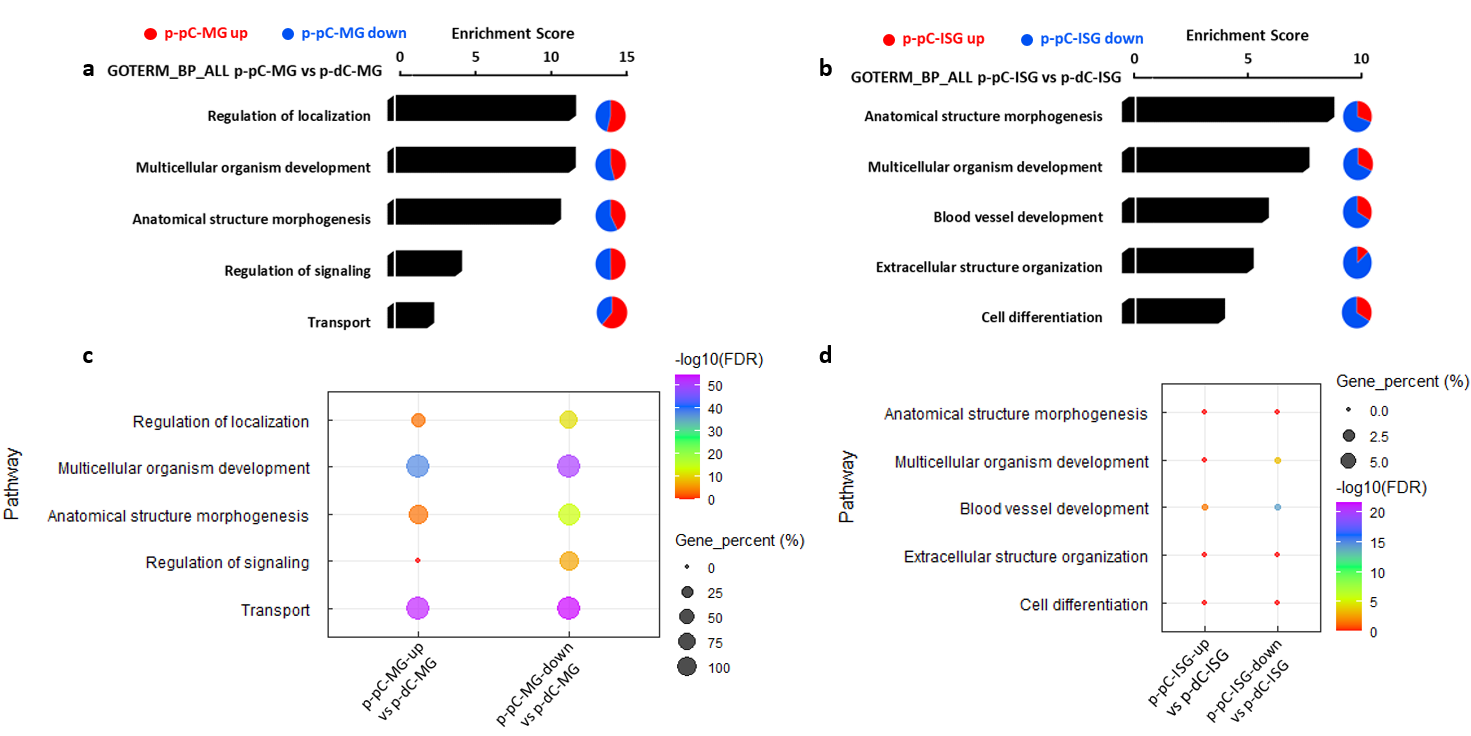

**Supplementary Fig. 2** Comparison of pathway enrichment in myenteric ganglia (MG) or inner submucosal ganglia (ISG) between porcine proximal and distal colon (p-pC, p-dC). a, b Enrichment of top five biological processes (BPs). The pie charts show the percentage of visible genes involved in the specified BPs. Red and blue color represents the genes with upregulated and downregulated expression levels in p-pC-MG or p-pC-ISG in comparison with p-dC-MG or p-dC-ISG, respectively. X-axis marks enrichment score, with the significance cut-off marked by the vertical white line (*p*-value < 0.05). c, d The bubble plots show the gene percentage and enrichment of the top BPs. Circle size represents the ratio of the number of pathway-specific differentially expressed genes (DEG) and the number of total DEG in each DEG list (Gene_percent). Color represents a -log10(FDR) distribution from big (orange) to small (purple). Up, upregulated expression. Down, downregulated expression.

**Supplementary Figure 3:**

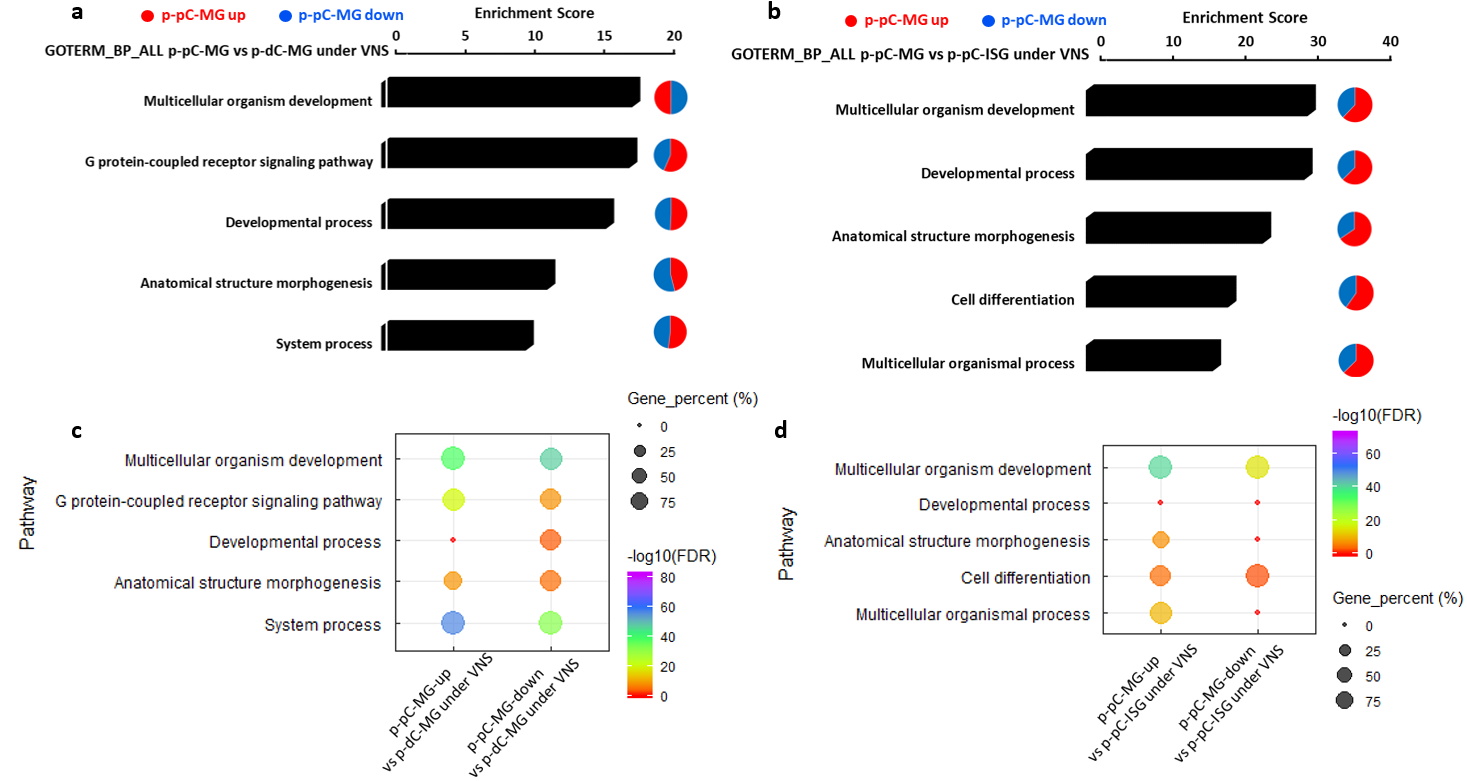

**Supplementary Fig. 3** Comparison of pathway enrichment in myenteric ganglia (MG) between porcine proximal and distal colon (p-pC, p-dC) and between p-pC-MG and inner submucosal ganglia (ISG) in p-pC with vagal nerve stimulation (VNS). a, b Enrichment of top five biological processes (BPs). The pie charts show the percentage of visible genes involved in the specified BPs. Red and blue color represents the genes with upregulated and downregulated expression levels in p-pC-MG in comparison with p-dC-MG or p-pC-ISG, respectively. X-axis marks enrichment score, with the significance cut-off marked by the vertical white line (*p*-value < 0.05). c, d The bubble plots show the gene percentage and enrichment of the top BPs. Circle size represents the ratio of the number of pathway-specific differentially expressed genes (DEG) and the number of total DEG in each DEG list (Gene_percent). Color represents a -log10(FDR) distribution from big (orange) to small (purple). Up, upregulated expression. Down, downregulated expression.

**Supplementary Figure 4:**

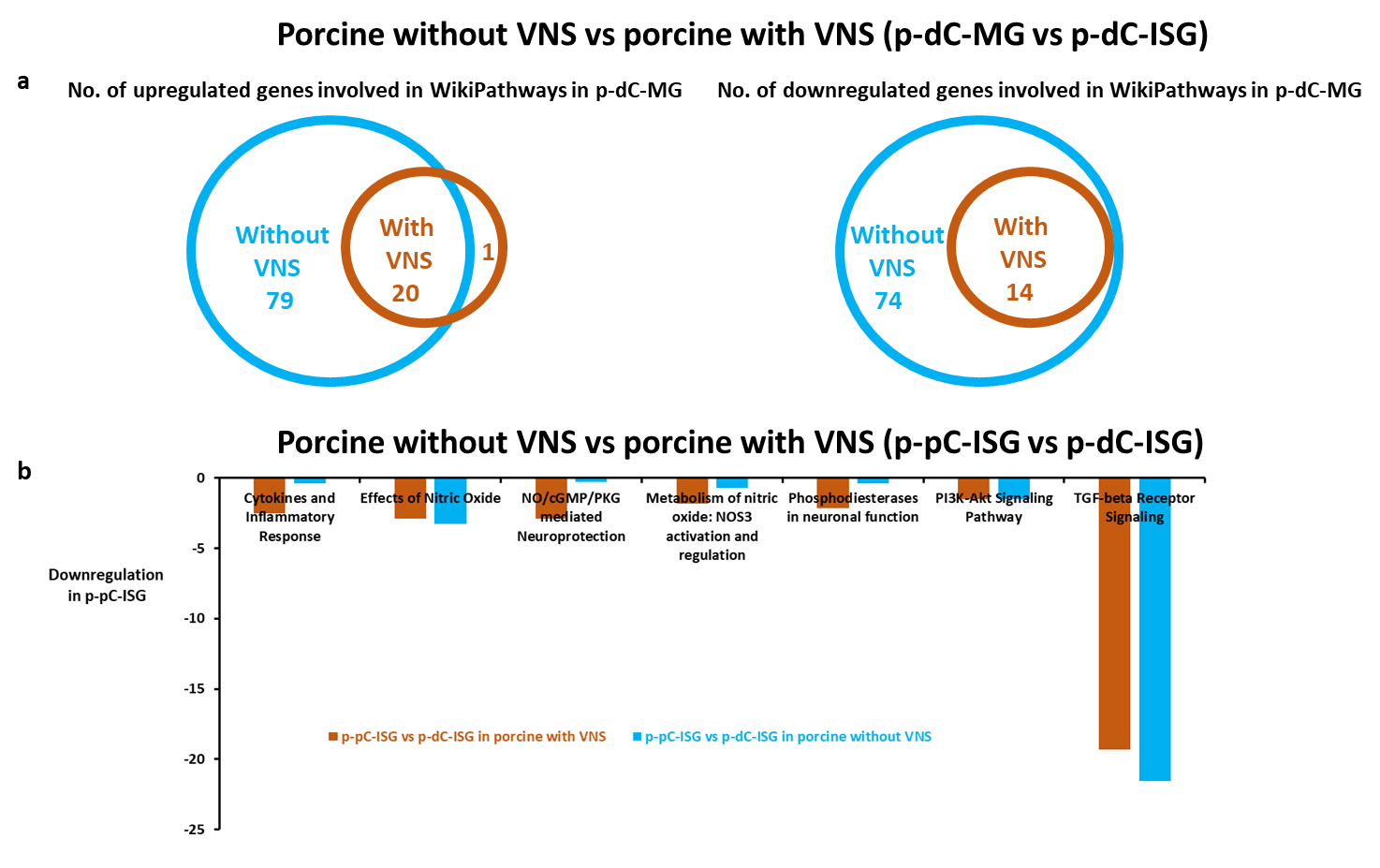

**Supplementary Fig. 4** Comparison of pathway enrichment between myenteric ganglia (MG) and inner submucosal ganglia (ISG) in porcine distal colon (p-dC) (a) and between ISG in proximal colon (p-pC) and p-dC-ISG (b) with and without vagal nerve stimulation (VNS). The Venn diagram illustrates the numbers of the differentially expressed genes involved in WikiPathways (a) and the bar graph shows the difference in WikiPathways enrichment (b). The light blue color represents the difference without VNS and the orange color represents the difference with VNS. Fig. b shows the enrichment ratio of the same WikiPathway involving upregulated and downregulated genes in each list of differentially expressed genes based on the bulk RNA sequencing data. The light blue bars represent the enrichment ratio without VNS and the orange bars represent the enrichment ratio with VNS.

**Supplementary Figure 5:**

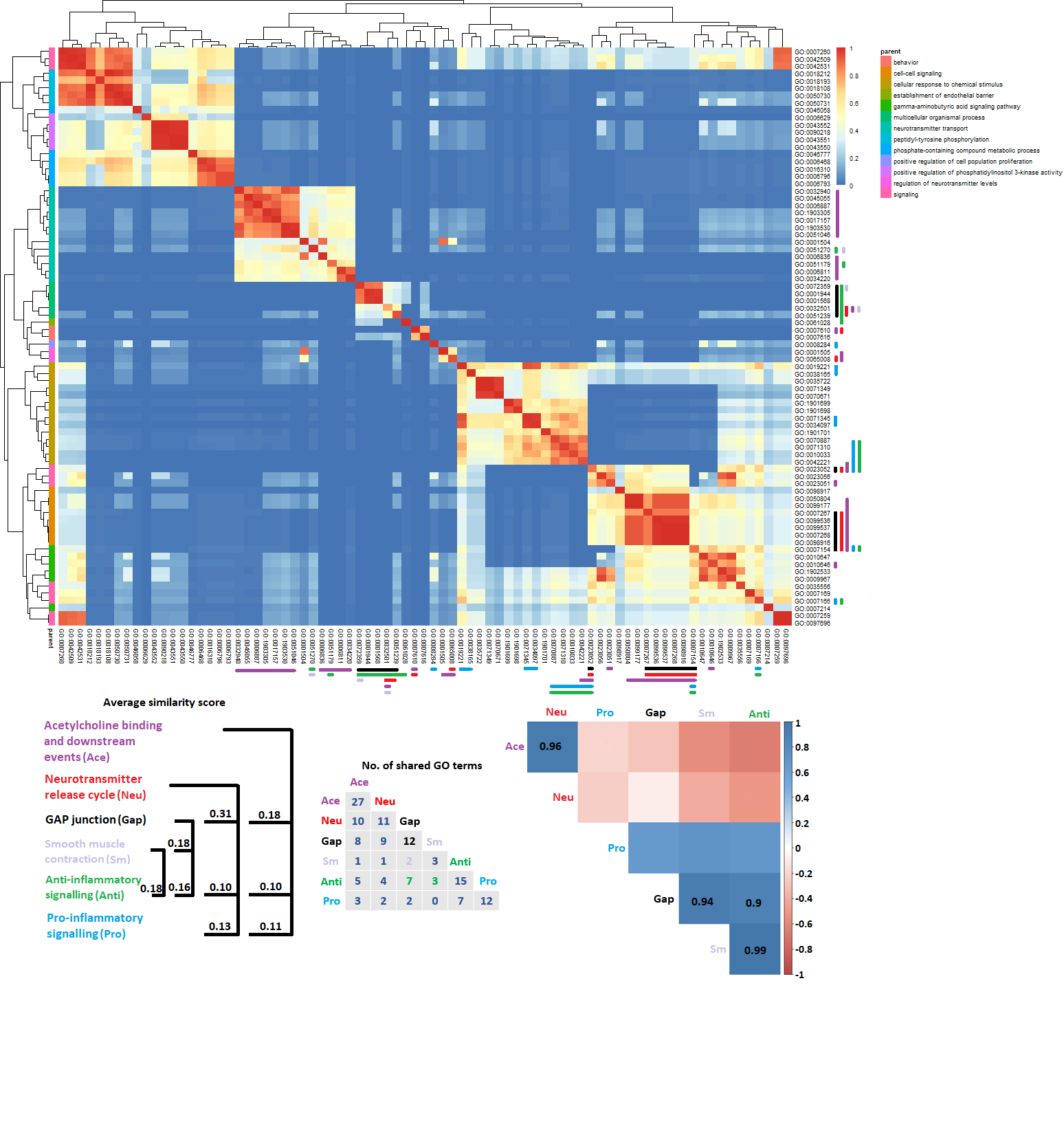

**Supplementary Fig. 5** Heatmap represents the similarity matrix between Gene Ontology (GO) terms in comparison of myenteric ganglia (MG) and inner submucosal ganglia (ISG) in porcine proximal colon (p-pC) with vagal nerve stimulation (VNS). Only biological processes were selected for the analysis. The child GO terms were defined in a number of parent terms and WikiPathways of interest. The number of shared GO terms between WikiPathways of interest and average similarity score between the WikiPathways were also summarized. Pearson correlation analysis was performed based on pathway enrichment scores. The insignificant correlations were left blank. The color bars represent the specified child GO terms classified into designated clusters.

**Supplementary Figure 6:**

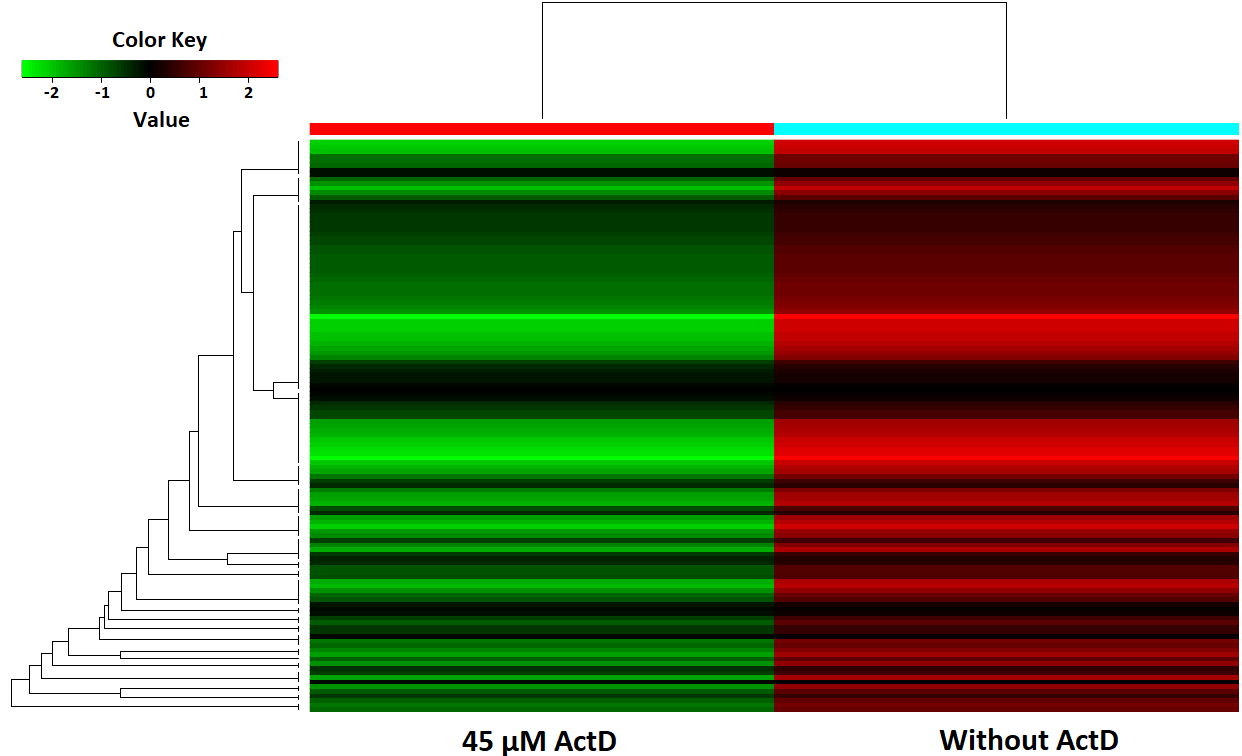

**Supplementary Fig. 6** Heatmap showing that addition of Actinomycin D (ActD) suppresses artificially induced expression levels of immediate-early genes (IEGs) during single-cell dissociation. After cell suspension preparation from muscularis externa of porcine proximal colon, more than 1 million cells per sample were collected for bulk RNA-seq. The expression levels of total 127 IEGs were extracted.

**Supplementary Table 1:**

**Receptors and ligands expressed in neuronal and glial subsets**

| **Receptor** | **Ligand** |
| --- | --- |
| AGTR1/Cholinergic neurons | VEGFA/ Nitrergic neurons |
| AGTR1/Cholinergic neurons | ALB/SLC41A1_Glia |
| ITGB1/Cholinergic neurons | VEGFA/ Nitrergic neurons |
| ITGB1/Cholinergic neurons | ALB/SLC41A1_Glia |
| ITGB1/Cholinergic neurons | SPP1/SLC41A1_Glia |
| ITGB1/Cholinergic neurons | TGFB3/SLC41A1_Glia |
| LDLR/Cholinergic neurons | CLU/SPC24_Glia |
| LDLR/Cholinergic neurons | ALB/SLC41A1_Glia |
| VCL/Cholinergic neurons | ALB/SLC41A1_Glia |
| GPC4/Glutamatergic neurons | PCOLCE2/ Cholinergic neurons |
| LEPR/Glutamatergic neurons | IL7/SLC41A1_Glia |
| LEPR/Glutamatergic neurons | IGF1/SLC41A1_Glia |
| TGFBR1/Glutamatergic neurons | TGFB3/SLC41A1_Glia |
| ACVR2B/ Nitrergic neurons | TGFB3/SLC41A1_Glia |
| ACVR2B/ Nitrergic neurons | IGF1/SLC41A1_Glia |
| GNG8/ Nitrergic neurons | C5/SLC41A1_Glia |
| ANPEP/SPC24_Glia | ALB/SLC41A1_Glia |
| ANPEP/SPC24_Glia | VEGFA/ Nitrergic neurons |
| CALCRL/SLC41A1_Glia | VEGFA/ Nitrergic neurons |

**Supplementary Table 2**

**Coverage of functional linkages involving top five WikiPathways in the investigated cell types**

| **Comparisons** | **Key DEG involved in the** **top five WikiPathways in the investigated cell types** | **Proportion of functional linkages involving the key DEG to those involving all DEG (%)** |
| --- | --- | --- |
| **p-pC-MG up vs p-dC-MG in porcine without VNS** | *MAPK3*, *PRKX*, *LDLR* | 96.16 |
| **p-pC-MG down vs p-dC-MG in porcine without VNS** | *GNG8*, *CXCL12*, *GUCY1A1*, *TGFBR3* | 99.13 |
| **p-pC-MG up vs p-dC-MG in porcine with VNS** | *CLTB*, *DAB2*, *MAPK3*, *NFKBIB*, *LYN*, *BCAR1*, *GRK5*, *PIK3CB*, *PRKX*, *GRK6*, *PTK2B*, *STAT1*, *EGR1*, *OSMR*, *LDLR*, *VEGFA*, *IL6ST*, *JAK1*, *ENG*, *TGIF1* | 99.76 |
| **p-pC-MG down vs p-dC-MG in porcine with VNS** | *TJP1*, *CXCL12*, *PIK3R1*, *TGFBR3*, *BMP4* | 98.27 |

**Note:** p-pC-MG and p-dC-MG, myenteric ganglia in porcine proximal and distal colon, respectively; VNS, vagal nerve stimulation; up, upregulated expression; down, downregulated expression; DEG, differentially expressed gene.

**Supplementary Table 3**

**Categories for the WikiPathways of interest**

| **Synaptic plasticity** | **Neurotransmitter binding** | | **Interactions between immune cells and neurons** | | **Neuroprotection** | | **Nitric Oxide** | | **Neuroinflammation** | | **Anti-inflammation** | | **Cytokine production** | | **Neurogenesis** |
| --- | --- | --- | --- | --- | --- | --- | --- | --- | --- | --- | --- | --- | --- | --- | --- |
| Synaptic Vesicle Pathway | Synaptic Vesicle Pathway | Oncostatin M Signaling Pathway | | NO/cGMP/PKG mediated Neuroprotection | | Effects of Nitric Oxide | |  | | TGF-beta Receptor Signaling | | Chemokine signaling pathway | | Dopaminergic Neurogenesis | |
| Glutamate binding, activation of AMPA receptors and synaptic plasticity | Glutamate binding, activation of AMPA receptors and synaptic plasticity | Cytokines and Inflammatory Response | |  | | Metabolism of nitric oxide: NOS3 activation and regulation | |  | |  | |  | |  | |
| Neurotransmitter receptors and postsynaptic signal transmission | Neurotransmitter receptors and postsynaptic signal transmission | COVID-19 Adverse Outcome Pathway | |  | | PI3K-Akt Signaling Pathway | | PI3K-Akt Signaling Pathway | |  | | PI3K-Akt Signaling Pathway | |  | |
| Neurotransmitter release cycle | Neurotransmitter release cycle |  | |  | |  | |  | |  | |  | |  | |
| Phosphodiesterases in neuronal function | Phosphodiesterases in neuronal function |  | |  | | Phosphodiesterases in neuronal function | |  | |  | |  | |  | |
| GABA receptor Signaling | GABA receptor Signaling |  | |  | |  | |  | |  | |  | |  | |
| Acetylcholine synthesis | Neurotransmitter uptake and metabolism In glial cells | Neurotransmitter uptake and metabolism In glial cells | |  | |  | |  | |  | |  | |  | |
| Cannabinoid receptor signaling | Acetylcholine binding and downstream events | Cannabinoid receptor signaling | |  | |  | |  | |  | |  | |  | |
| Nicotine Activity on Dopaminergic Neurons | Neurotransmitter clearance | Nicotine Activity on Dopaminergic Neurons | |  | |  | |  | |  | |  | |  | |

**Supplementary Table 4**

**Well-known cytokines and the receptors used for the analyses in our study**

| **Pro-inflammatory signaling** | |
| --- | --- |
| **IL-1** | IL1A, IL1B, IL18, IL33, IL36A, IL36B |
| **IL-6** | IL11, IL6R, CNTFR, CTF1, LIF, SPP1, OSM, OSMR |
| **TNFα** | TNF, LTA, TNFSF13B, TNFSF13 |
| **IL-17** | IL17A, IL17B, IL17C, IL17D, IL17F, IL25 |
| **IFN** | IFNB1, IFNAR1, IFNK, IFNL1 |
| **C-C Motif chemokine** | CCR2, CCL2 |
| **C-X-C Motif chemokine** | CXCL12 |
| **Anti-inflammatory signaling** | |
| **IL-12** | IL12RB1, IL12RB2, IL23R, IL23A, IL27 |
| **IL-10** | IL10, IL19, IL20, IL24, IL22, IL26 |
| **TGFβ** | TGFB1, TGFBR1, TGFBR2, TGFBR3 |
| **C-X-C Motif chemokine** | CXCL11 |
